# Thalamic spindles and upstates, but not ripples, coordinate cortico-cortical and hippocampo-cortical co-ripples in humans

**DOI:** 10.1101/2022.09.15.507471

**Authors:** Charles W. Dickey, Ilya A. Verzhbinsky, Sophie Kajfez, Burke Q. Rosen, Christopher E. Gonzalez, Patrick Y. Chauvel, Sydney S. Cash, Sandipan Pati, Eric Halgren

## Abstract

The co-occurrence of brief ∼90Hz oscillations (co-ripples) may be important for integrating information across the neocortex and hippocampus and, therefore, essential for sleep consolidation, and cognition in general. However, how such co-ripples are synchronized is unknown. We tested if cortico-cortical and hippocampal-cortical ripple co-occurrences are due to the simultaneous direct propagation of thalamic ripples, and/or if they are coordinated by lower frequency thalamic waves. Using human intracranial recordings, we found that ripples are generated in the anterior and posterior thalamus during local spindles on the down-to-upstate transition in non-rapid eye movement sleep, with similar characteristics as cortical and hippocampal ripples. However, thalamic ripples only infrequently co-occur or phase-lock, with cortical and hippocampal ripples. In contrast, thalamo-cortical spindles and upstates were strongly coordinated with cortico-cortical and hippocampo-cortical co-rippling. Thus, while thalamic ripples may not directly drive multiple cortical or hippocampal sites at ripple frequency, thalamo-cortical spindles and upstates may provide the input necessary for spatially distributed co-rippling to integrate information in the cortex.

**Significance Statement:** Widespread networks of ∼90 Hz oscillations, called “ripples,” have recently been identified in humans and may help to bind information in the cortex and hippocampus for memory. However, it is not known whether the thalamus generates ripples, and if so whether they, or other thalamic waves, coordinate networks of co-occurring cortical and hippocampal ripples. Here, we show that the human thalamus generates ∼90 Hz ripples during NREM sleep. While thalamic ripples do not appear to directly synchronize ripple co-occurrence in the cortex and hippocampus, our data provide evidence that propagating thalamo-cortical spindles and upstates organize these networks. Thus, the thalamus projects slower frequency waves that modulate higher frequency hippocampo-cortical oscillatory networks for memory in humans.

## Introduction

Ripples are brief high-frequency oscillations of the local field potential (LFP). In rodent hippocampus during non-rapid eye movement sleep (NREM), ripples mark the replay of spatiotemporal firing patterns that are crucial in the consolidation of declarative memories (Wilson and McNaughton, 1994; Ego-Stengel and Wilson, 2009; Buzsáki, 2015; Maingret et al., 2016). In humans, ∼90Hz ripples are generated in the cortex during both NREM and wakefulness (Dickey et al., 2022a; Dickey et al., 2022b). Cortical ripples couple to cortical and hippocampal ripples preceding recall (Vaz et al., 2019; Dickey et al., 2022b), and entrain the replay of neuron firing sequences established during encoding (Vaz et al., 2020).

Spindles are 10-16Hz oscillations that occur during NREM and are important for memory consolidation (Adamantidis et al., 2019). They are generated through interactions of intrinsic currents and local circuits in the thalamus (Destexhe and Sejnowski, 2003; McCormick et al., 2015) and are projected to all cortical areas (Mak-McCully et al., 2017; Piantoni et al., 2017), often on the down-to-upstate transition (Gonzalez et al., 2018), timing that is important for memory consolidation (Klinzing et al., 2019). Downstates, where cortical neurons are silent, and upstates, where cortical neurons fire at near-waking levels, comprise the slow oscillation and K-complex of human NREM (Cash et al., 2009; Csercsa et al., 2010). Recurrent thalamo-cortical projections support the initiation, synchronization, and termination of spindles and down-to-upstates (Contreras et al., 1996; Mak-McCully et al., 2017; Crunelli et al., 2018). Critical to a possible role in consolidation, human cortical ripples occur during spindles just prior to the upstate peak (Dickey et al., 2022a), and may mark the replay of cell-firing encoding motor patterns learned in prior waking (Rubin et al., 2022). Furthermore, hippocampal ripples are coordinated with cortical ripples (Vaz et al., 2019; Vaz et al., 2020; Dickey et al., 2022b), spindles, downstates, and upstates (Jiang et al., 2019a, b).

Crucially, human cortical ripples co-occur and phase-lock across long distances in multiple lobes in both hemispheres (Dickey et al., 2022b). Neither co-occurrence nor phase-locking decreases with separation between rippling locations, implying either a network of densely connected coupled cortical oscillators, or a central subcortical structure that broadly triggers and synchronizes cortical ripples. The thalamus is a prime candidate for this role given its direct anatomical connections with both the hippocampus and cortex (Varela et al., 2014), as well as its ability to modulate and synchronize spindles and down-to-upstates across widespread cortical areas (Mak-McCully et al., 2014; Piantoni et al., 2016; Mak-McCully et al., 2017; Gonzalez et al., 2018). Consolidation during NREM in mice is enhanced by optogenetic stimulation of the thalamus that evokes cortical spindles co-occurring with cortical upstates and hippocampal sharpwave-ripples (Latchoumane et al., 2017). Thus, the thalamus could synchronize ripples, either directly by generating and widely projecting ripples, or indirectly by projecting spindles or upstates, which in turn evoke ripples in multiple cortical and hippocampal locations.

Ripples have not been previously studied in the human thalamus. We analyzed a rare collection of intracranial recordings from non-epileptogenic and non-lesioned anterior and posterior thalamus, with simultaneous recordings in the hippocampus, and cortex obtained from patients with epilepsy undergoing seizure focus localization. to determine if physiological ripples occur in the human thalamus, and if so, to elucidate their characteristics and relationships with cortical and hippocampal ripples. We subsequently tested if thalamic ripples co-occur and phase-lock with cortical or hippocampal ripples, as would be expected if thalamic ripples directly synchronize ripples in these regions. We found that the human thalamus generates ∼90Hz ripples during spindles on the down-to-upstate transition during NREM. Thalamic ripples were only weakly related to cortical or hippocampal ripples. However, the probability of co-rippling between cortical sites was greatly increased during thalamo-cortical spindles and upstates. Thus, the thalamus does not appear to drive widespread cortical ripple networks through the projection of ripples, but may instead coordinate ripple co-occurrence by synchronously modulating cortical sites through the projection of thalamic spindles and upstates.

## Results

### Thalamic ripples are generated during human NREM sleep

We detected ripples in the thalamus, cortex, and hippocampus of 13 patients undergoing seizure focus localization with intracranial electrodes (Supp. Table 1-2; Supp. Fig.1-2). All channels included in the study were trans-gray matter bipolar derivations to minimize effects due to volume conduction, and localized to non-lesioned, non-epileptogenic tissue. Electrodes were implanted in anterior thalamic nuclei, projecting mainly to lateral, medial and orbital prefrontal cortices, and the cingulate gyrus (Supp. Table 1). Electrodes were also implanted in posterior thalamic nuclei, projecting mainly to posterior cortical areas. Sleep staging was performed to select NREM. Channels and epochs were only included if they did not contain frequent interictal spikes or artifacts. Ripples were detected using a previously published method (Dickey et al., 2022a; Dickey et al., 2022b), requiring increased amplitude and at least 3 distinct oscillation cycles within 70-100Hz. Putative ripples were rejected if they contained possible interictal spikes or artifacts.

We found that in addition to the cortex and hippocampus, ripples were generated in the human anterior and posterior thalamus during NREM (Fig.1A-D). These ripples were centered focally around ∼90Hz with a preceding increase in delta (0.1-4Hz) and concurrent increase in spindle (10-16Hz) band power. Thalamic ripple detections did not appear to be due to volume conduction since adjacent non-thalamic bipolar channels in the white matter did not demonstrate ripples at the times of the thalamic ripples (Supp. Fig.3). Furthermore, ripples were very infrequently detected in adjacent non-thalamic bipolar channels in the white matter (Supp. Fig.4).

**Figure 1.**
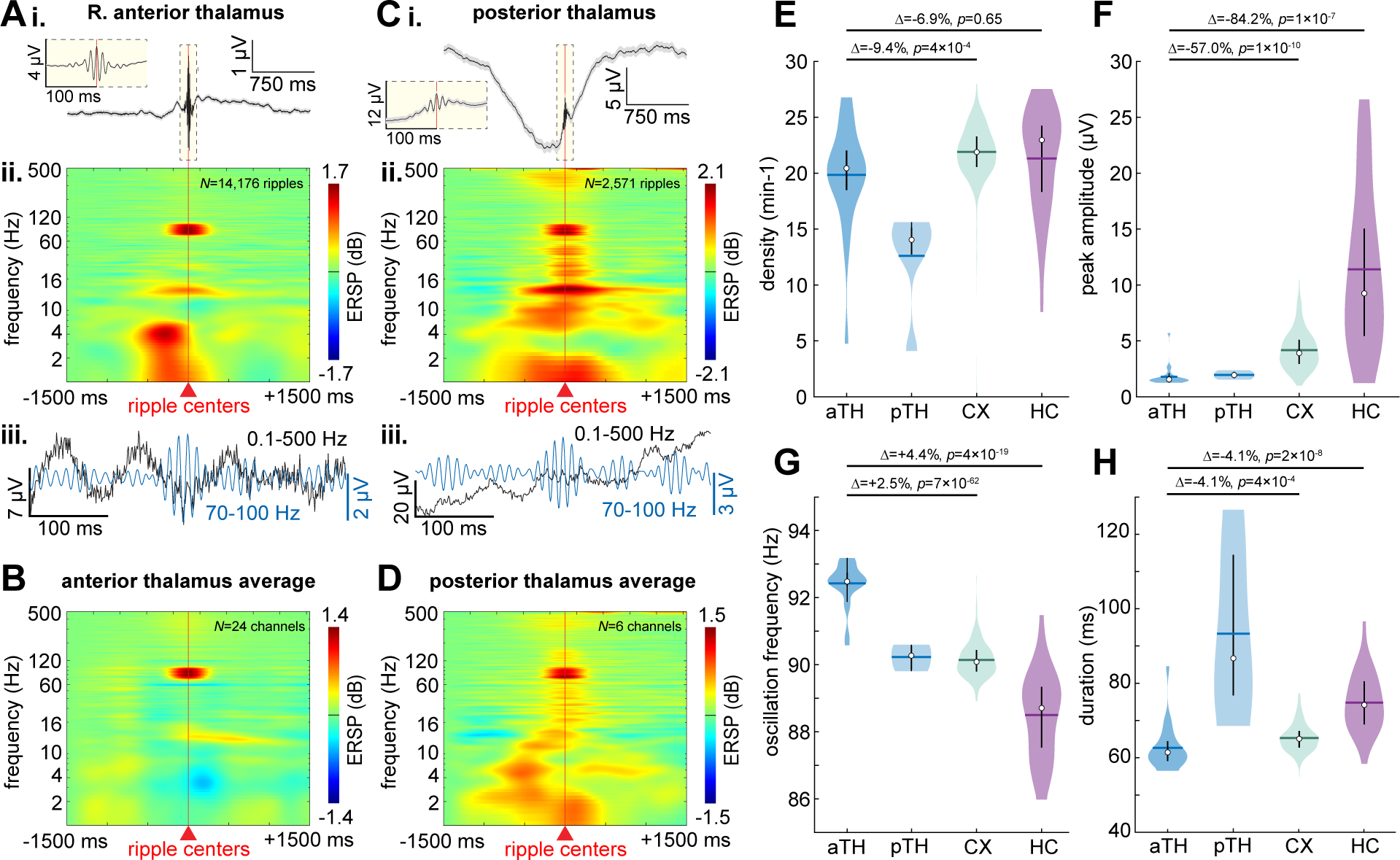
Ripples are generated in the human thalamus during NREM with similar characteristics as cortical and hippocampal ripples. (**A**) Ripples in one anterior thalamic bipolar channel: (**i**) average and SEM broadband LFP, (**ii**) average time-frequency, and (**iii**) example single sweep broadband (black) and 70-100Hz bandpass (blue). In the time-frequency plot, note the focally increased power at ∼90Hz, concurrent with increased ∼12Hz spindle band power, shortly following increased ∼4Hz delta power. In the single sweep, note the ripple occurrence on a ∼12Hz spindle. (**B**) Grand channel average time-frequency of all anterior thalamic channels (*N*=24 channels from patients 1-10). (**C-D**) Same as A-B except posterior thalamic ripples (*N*=6 channels from patients 11-13). (**E-H**) Anterior thalamic, posterior thalamic, cortical, and hippocampal ripple densities (**E**), peak 70-100Hz analytic amplitudes (**F**), oscillation frequencies (**G**), and durations (**H**) during NREM across all channels (*N_aTH_*=24, *N*_pTH_=6, *N_CX_*=261, *N_HC_*=31 channels). Note the consistent oscillation frequencies of ∼90Hz. Horizontal lines, means; circles, medians; vertical lines, interquartile ranges. Statistics performed with linear mixed-effects models with patient as random effect. *P*-values FDR-corrected for multiple comparisons across all channels from all patients. Distributions across individual ripples are shown in Supp. Fig. 6. aTH, anterior thalamus; CX, cortex; FDR, false discovery rate; HC, hippocampus; LFP, local field potential; NREM, non-rapid eye movement sleep; pTH, posterior thalamus; SEM, standard error of the mean.

### Thalamic ripples have similar characteristics as hippocampal and cortical ripples

Anterior thalamic ripples during NREM had a mean and standard deviation density (occurrence rate) of 19.8±4.6min^−1^, peak 70-100Hz analytic amplitude of 1.79±0.91µV, oscillation frequency of 92.4±0.6Hz, and duration of 62.6±5.9ms (Fig.1E-H; Supp. Table 3). Posterior thalamic ripples had a density of 11.3±2.2min^−1^, amplitude of 1.95±0.34µV, frequency of 90.2±0.4Hz, and duration of 93.3±23.0ms. Cortical ripples had an average density of 21.9±2.5min^−1^, amplitude of 4.17±1.69µV, frequency of 90.1±0.5Hz, and duration of 65.3±3.6ms (Supp. Fig.5 shows cortical ripples from patients with posterior thalamic leads). Hippocampal ripples had an average density of 21.3±4.7min^−1^, amplitude of 11.39±7.23µV, frequency of 88.5±1.3Hz, and duration of 74.8±7.9ms. Overall, these characteristics were consistent when analyzed across individual ripples (Supp. Fig.6) and across channels for individual patients (Supp. Fig.7). While thalamic ripples had smaller amplitudes than cortical or hippocampal ripples, they were easily distinguished from the baseline signal (Fig.1Aiii,Ciii), which was also smaller. These characteristics were similar to those of hippocampal and cortical ripples, with the exception of hippocampal ripples having larger amplitudes (Fig.1E-H; Supp. Table 3). Posterior thalamic ripples had a slightly lower density, similar amplitudes, slightly lower frequencies, and longer durations than anterior thalamic ripples. The relative sizes of ripples in thalamus, cortex, and hippocampus we found are consistent with the degrees of lamination of neurons and synaptic inputs in the three structures, as well as previous observations that human ripples are larger in hippocampus than cortex (Dickey et al., 2022a), and that spindles and downstates are larger in cortex than thalamus (Mak-McCully et al., 2017).

### Thalamic ripples are generated during local spindles on the down-to-upstate transition

It is thought that the downstate is initiated in the cortex and then projects to the thalamus, where hyperpolarization releases h and T currents leading to the generation of a thalamic spindle that is then projected back to the cortex at the time of the down-to-upstate transition (Mak-McCully et al., 2017). This precisely timed sequence is important for memory consolidation. We found that, on average, anterior and posterior thalamic ripples occur approximately 225-275ms following thalamic downstate peaks (Fig.2A-D; *N*=9/24 anterior thalamic channel pairs with significant modulations, post-FDR *p*<0.05, randomization test; significance across all channel pairs within −500-0 ms: *p*=2×10^−19^, *z*=8.9, and 0-500 ms: *p*=0.47, *z*=-6×10^−2^, one-sided Wilcoxon signed-rank test, Supp. Fig.8A; *N*=2/6 posterior thalamic, −500-0 ms: *p*=6×10^−5^, *z*=3.9, and 0-500 ms: *p*=0.99, *z*=2.2, Supp. Fig.8B). Of the 9 anterior thalamic channels with significant modulations, 7 had a significant order preference, all with downstates preceding ripples (post-FDR *p*<0.05, binomial test, expected value=0.5). Similarly, of the 2 posterior thalamic channels with significant modulations, both had a significant order preference with downstates preceding ripples. Anterior thalamic ripples tended to occur during thalamic spindles, shortly following their onsets (Fig.2E; *N*=7/24, −500 to 0 ms: *p*=1×10^−16^, *z*=8.2, and 0-500 ms: *p*=2×10^−12^, *z*=6.9, Supp. Fig.8C). Posterior thalamic ripples also coupled to thalamic spindles, albeit with a greater delay (∼350ms) following spindle onset (Fig.2F; *N*=1/6, −500-0 ms: *p*=3×10^−4^, *z*=3.4, and 0-500 ms: *p*=0.92, *z*=-1.4, Supp. Fig.8D). Anterior and posterior thalamic ripples occurred ∼125s and ∼275ms, respectively, preceding thalamic upstate peaks (Fig.2G-H; anterior thalamus *N*=12/24, −500-0 ms: *p*=0.97, *z*=1.92, and 0-500 ms: *p*=9×10^−17^, *z*=8.24, Supp. Fig.8E; posterior thalamus *N*=1/6, −500-0 ms: *p*=0.028, *z*=-1.9, and 0-500 ms: *p*=0.09, *z*=1.3, Supp. Fig.8F). Of the 12 anterior thalamic channels with significant modulations, 9 had a significant order preference, all with thalamic ripples preceding thalamic upstate peaks. Results from all channels confirm these relationships (Supp. Fig.8). Overall, these data show that thalamic ripples tend to occur on local down-to-upstates (Fig.2I-J), especially during spindles (Fig.K-L).

**Figure 2.**
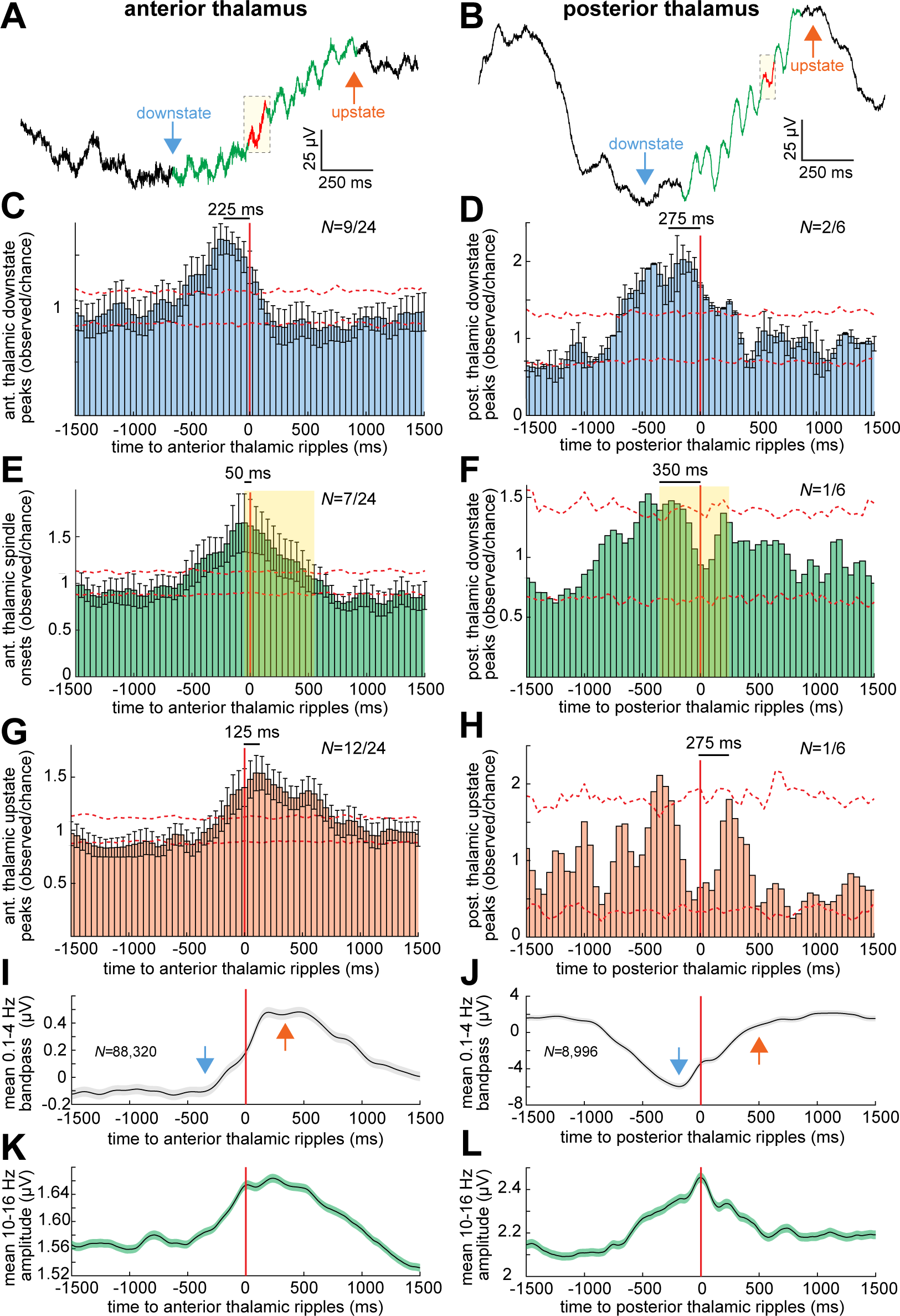
Thalamic ripples occur on the local down-to-upstate transition during spindles, similar to cortical ripples. (**A-B**) Example broadband single sweep anterior (**A**) and posterior (**B**) thalamic ripple occurring during a spindle on the down-to-upstate transition. (**C-D**) Average and SEM times of anterior (**C**) and posterior (**D**) thalamic downstate peaks relative to thalamic ripples at *t*=0 across channels that showed significant coupling in order to demonstrate effect time courses (*N_aTH_*=9/24 and *N_pTH_*=2/6 significant channels, post-FDR *p*<0.05, randomization test). See Supp. Fig.8 for results across all channels. (**E-F**) Same as C-D except thalamic spindle onsets (*N_aTH_*=7/24 and *N_pTH_*=1/6). Shaded box denotes the average spindle interval. (**G-H**) Same as C-D except thalamic upstate peaks (*N_aTH_*=12/24, *N_pTH_*=1/6). (**I-J**) Average and SEM anterior (**I**) and posterior (**J**) thalamic ripple-locked 0.1-4Hz delta bandpass (*N_aTH_*=88,320 and *N_pTH_*=8,996 ripples). (**K-L**) Same as I-J except 10-16Hz spindle band analytic amplitude. Dashed error shows 98% confidence interval of the null distribution.

### Thalamic ripples infrequently and weakly couple to and co-occur with cortical and hippocampal ripples

Previously, we showed that ripples frequently and strongly co-occur between cortical sites (i.e., cortico-cortical co-ripples) in widespread networks, and also (less strongly) between the hippocampus and cortex (i.e., hippocampo-cortical co-ripples) (Dickey et al., 2022b). One possible mechanism underlying this is a thalamo-cortical driving circuit. Therefore, we tested if thalamic ripples couple (within ±250ms) or co-occur (“co-ripple”; ≥25ms overlap) with cortical or hippocampal ripples. Confirming our previous results in a different dataset, we found frequent (>50% of channel pairs) and strong (>25% peak increase in probability from chance) cortico-cortical ripple coupling (Fig.3A-B). By contrast, anterior and posterior thalamic ripples infrequently (<50% of channel pairs significant) and weakly (<25% peak increase in probability from chance) coupled with cortical ripples (Fig.3C-D). Replicating our previous findings, hippocampo-cortical ripple coupling was frequent but weak (Fig.3E). By contrast, there was infrequent and weak coupling between anterior thalamic and hippocampal ripples (Fig.3F). Thalamic ripple auto-correlation did not reveal rhythmic entrainment by downstates or upstates (Supp. Fig.9). Results from all channel pairs confirm these relationships (Supp. Fig.10). Notably, there was a significantly increased probability of cortico-cortical co-rippling given rippling on a different randomly selected cortical channel compared to the probability of rippling on a randomly selected anterior thalamic channel (mean±SEM: 0.18±0.003 vs. 0.09±0.001%; *p*=1×10^−233^, *t*(3897)=34.8; one-sided paired *t*-test) as well as posterior thalamic rippling on a randomly selected channel (mean±SEM: 0.67±0.09 vs. 0.17±0.03%; *p*=6×10^−7^, *t*(27)=6.2.

**Figure 3.**
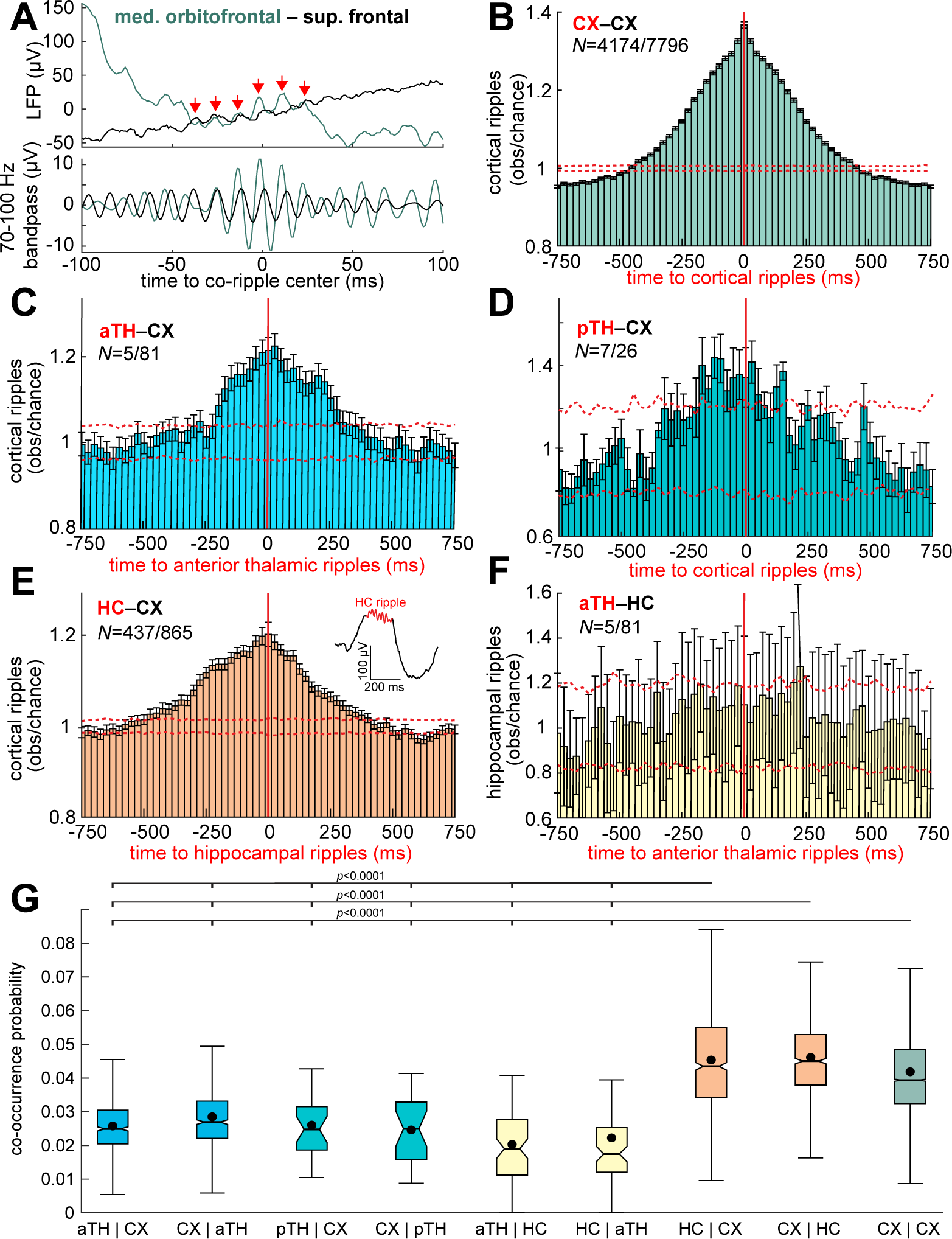
Thalamic ripples infrequently and weakly co-occur with cortical and hippocampal ripples during NREM. (**A**) Example co-occurring thalamo-cortical, cortico-cortical, and hippocampo-cortical ripples. (**B**) Average observed over chance cortical ripple centers in one channel relative to cortical ripple centers in another channel for significant channel pairs (*N*=4174/7796). Note that no cortical bipolar channels included shared a common contact. Error bars show SEM and dashed error shows 98% confidence interval of the null distribution. See Supp. Fig.10 for results from all channel pairs. (**C**) Cortical ripples relative to anterior thalamic ripples (*N*=88/649 significant, post-FDR *p*<0.05, randomization test; 25ms Gaussian smoothed with σ=5ms). (**D**) Same as C except for posterior thalamic ripples (*N*=7/26). (**E**) Same as B except hippocampal ripples relative to cortical ripples (*N*=437/865). Note the single sweep example of a hippocampal ripple on a sharpwave. (**F**) Same as B except hippocampal ripples relative to anterior thalamic ripples (*N*=5/81). (**G**) Conditional probabilities of ripple co-occurrences (≥25ms overlap) between channels (e.g., aTH | CX = probability of an anterior thalamic ripple co-occurring with a given cortical ripple). Post-FDR *p*-values, two-sided two-sample *t*-test. Error bars show SEM.

Cortico-cortical and hippocampo-cortical sites also were more likely to co-ripple than thalamo-cortical and thalamo-hippocampal (Fig.3G). Among anterior thalamo-cortical channel pairs, 14% (88/649) were significantly coupled (post-FDR *p*<0.05, randomization test) and 6% (40/649) had a significant number of co-occurrences above chance (post-FDR *p*<0.05, randomization test). Among posterior thalamo-cortical channel pairs, 27% (7/26) were significantly coupled and 31% (8/26) significantly co-occurred. Among anterior thalamo-hippocampal pairs, even fewer, only 6% (5/81) were significantly coupled and 2% (2/81) had a significant number of co-occurrences. In contrast, among cortico-cortical pairs (including both permutations for each pair), 54% (4174/7796) were significantly coupled and 47% (1828/3898) had a significant number of co-occurrences. Among hippocampo-cortical pairs, 51% (437/865) were significantly coupled and 15% (126/865) had significant co-occurrences. All of these coupling relationships except thalamo-hippocampal were significant across all channels from all patients (cortico-cortical: *p*<1×10^−25^, *z*=257.1, one-sided Wilcoxon signed-rank test; anterior thalamo-cortical: *p*<1×10^−25^, *z*=40.1; posterior thalamo-cortical: *p*=6×10^−23^, *z*=9.8; hippocampo-cortical: *p*<1×10^−25^; anterior thalamo-hippocampal *p*=0.58, *z*=-0.2. Anterior thalamo-cortical co-ripple coupling was significantly less common than cortico-cortical (*p*<0.00001, χ^2^=383.1, *df*=1) or hippocampo-cortical (*p*<0.00001, χ^2^=88.1, *df*=1), which was also the case for co-occurrences (thalamo-cortical vs. cortico-cortical: *p*<0.00001, χ^2^=381.4, *df*=1; thalamo-cortical vs. hippocampo-cortical: *p*<0.00001, χ^2^=26.8, *df*=1). Across these four types of region pairs, the proportion of channel pairs with significant ripple coupling was significantly different (*p*<0.00001, χ^2^=438.73, *df*=3), as was the proportion of channel pairs with significant numbers of ripple co-occurrences above chance (*p*<0.00001, χ^2^=485.3, *df*=3).

### Thalamic ripples do not phase-lock with cortical or hippocampal ripples

We previously showed that cortico-cortical but not hippocampo-cortical co-ripples often strongly phase-lock, suggesting a possible intracortical mechanism (Dickey et al., 2022b). However, it is also possible that phase-locking between cortical locations is driven by a common input from thalamic ripples. To determine if channel pairs had phase-locked ripples, we computed phase-locking values (PLVs) (Lachaux et al., 1999), which measure phase consistency independent of amplitude, here using the 70-100Hz phase across co-occurring ripples (≥25ms overlap) at each time point for each channel pair (≥40 co-ripples required per pair). We found only one thalamo-cortical site pair had significantly phase-locked co-ripples (Supp. Fig.11A; *N*=1/451 significant channel pairs, post-FDR *p*<0.05, randomization test), and no thalamo-hippocampal site pairs had significantly phase-locked co-ripples (Supp. Fig.11B). Consistent with our previous results, cortico-cortical pairs had significantly phase-locked co-ripples (Supp. Fig.11C; *N*=52/2833), specifically about 8x as many as thalamo-cortical pairs, and no hippocampo-cortical pairs had phase-locked co-ripples (Supp. Fig.11E; *N*=0/490), nor did any thalamo-hippocampal pairs (Supp. Fig.11F; *N*=0/47. Across these 4 types of region pairs, the proportion of phase-locked channel pairs was significantly different (*p*=0.001, χ^2^=16.2, *df*=3). Thus, these results support our hypothesis that networks of phase-locked cortico-cortical co-ripples are directly driven intracortically rather than from thalamic or hippocampal inputs.

### Thalamo-cortical and thalamo-hippocampal co-spindles phase-lock

Given that cortical ripples occur during cortical spindles and phase-lock between cortical sites and between the cortex and hippocampus, we hypothesized that the thalamus organizes these networks by projecting spindles and upstates to the cortex (Fig.4A). To test this hypothesis, we first computed 10-16Hz PLVs between spindles in the thalamus and cortex. Since the anterior thalamus has more projections to the anterior than posterior cortex, we analyzed these regions separately (Fig.4B). In our analysis, the anterior cortex included orbitofrontal, prefrontal, and cingulate cortices, whereas the posterior cortex included all other cortical areas. We found significant thalamo-cortical spindle PLV modulations between anterior thalamus and anterior cortex (Fig.4C; *N*=188/369 significant pairs, post-FDR *p*<0.05, randomization test) as well as between anterior thalamus and posterior cortex (*N*=71/280). There was a significantly greater proportion of significant thalamo-cortical channel pairs between anterior thalamus and anterior cortex compared to anterior thalamus and posterior cortex (*p*<0.00001, χ^2^=43.5, *df*=1). There was no significant thalamo-cortical spindle PLV between posterior thalamus and anterior cortex (Fig.4D; *N*=0/4), but there was significant thalamo-cortical spindle PLV between posterior thalamus and posterior cortex (*N*=9/22). Furthermore, there was a greater mean PLV modulation amplitude within ±250ms around the thalamo-cortical temporal centers in the anterior compared to posterior cortex (*p*=5×10^−15^, *t*(613)=7.9, one-sided two-sample *t*-test). Results by individual cortical parcel are shown in Supp. Fig.12 These results are consistent with preferential projections of the anterior thalamus to the anterior cortex and the posterior thalamus to the posterior cortex. Notably, our data show prominent phase-locking between thalamo-cortical spindles but not ripples (Fig.4C-E; Supp. Fig.11A-B). We also evaluated 10-16 Hz phase-locking of spindles co-occurring between the anterior thalamus and hippocampus and found that 31/81 (38%) thalamo-hippocampal channel pairs had significantly phase-locked co-spindles.

**Figure 4.**
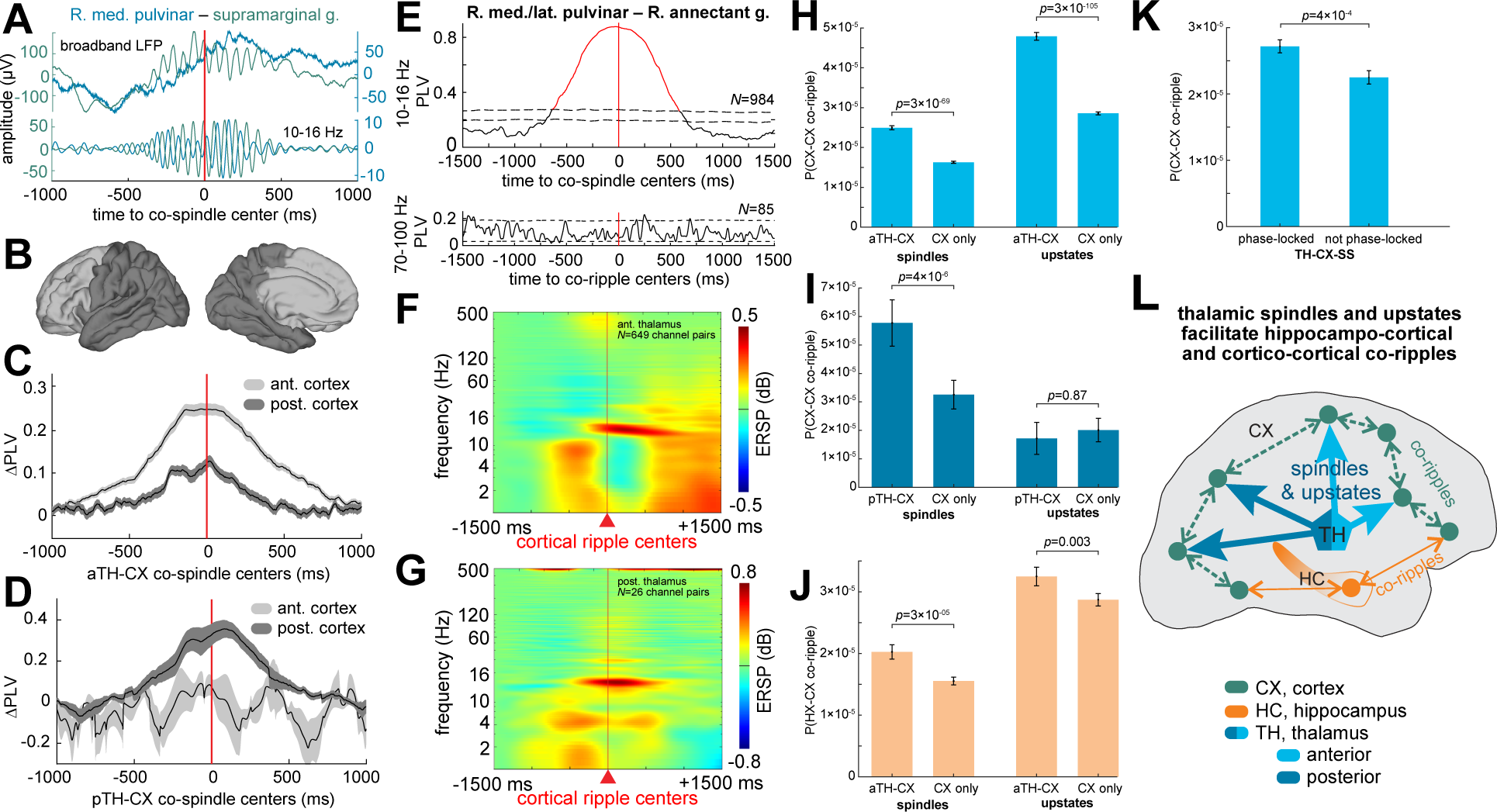
Cortico-cortical and hippocampo-cortical co-rippling are enhanced during thalamo-cortical spindles and upstates. (**A**) Example thalamo-cortical co-spindle in broadband and 10-16Hz bandpass. (**B**) Division of the cortex into anterior and posterior regions based on connectivity with the anterior and posterior thalamus. (**C-D**) Mean and standard deviation 10-16Hz PLVs of thalamo-cortical co-spindles, including anterior thalamus (**C**) and posterior thalamus (**D**), across channel pairs in anterior cortex (*N_aTH_*=188/369 and *N_pTH_*=9/22 significant, post-FDR *p*<0.05, randomization test) compared to posterior cortex (*N_aTH_*=71/280 and *N_pTH_*=0/4). Note greater phase-locking between anterior thalamus with anterior cortex, and for posterior thalamus with posterior cortex. Results by cortical parcel are shown in Supp. Fig.12. (**E**) Example thalamo-cortical pair showing very high 10-16Hz co-spindle phase-locking with no detectable 70-100Hz co-ripple phase-locking. (**F-G**) Average time-frequency of anterior (**F**) and posterior (**G**) thalamus time-locked to cortical ripples (*N*_aTH_=649 and *N*_pTH_=26). (**H**) Cortico-cortical co-ripple probability across cortico-cortical channel pairs when the anterior thalamus is spindling with either cortical channel versus when it is not (left bars; *p*=3×10^−69^, *t*(2915)=18.0; one-sided paired *t*-test), and similarly when the anterior thalamus and cortex share upstates versus when they do not (right bars; *p*=3×10^−105^, *t*(2948)=22.7). (**I**) Same as H except posterior thalamus (spindles: *p*=4×10^−6^, *<u>t</u>*<u>(17)</u>=6.2; upstates: *p*=0.87; *t*(27)=1.1). (**J**) Same as H except hippocampo-cortical co-ripple probability (spindles: *p*=3×10^−5^, *t*(544)=4.0; upstates: *p*=0.003, *t*(544)=2.7). (**K**) Same as H except cortico-cortical co-ripple probability when one versus neither cortical channel has 10-16Hz phase-locked spindles with the thalamus (in both cases TH and at least one of the CX channels are co-spindling; *p*=4×10^−4^, *t*(1936)=3.3). (**L**) Proposed synergistic mechanisms for achieving widely distributed cortical ripples that co-occur and phase-lock. Cortical depolarization via thalamo-cortical spindles and upstates (blue) trigger ripples which phase-lock through cortico-cortical coupling (green), and co-occur with hippocampal ripples via direct connections (orange). Our data do not support the alternative hypothesis wherein thalamic ripples directly drive cortical and/or hippocampal ripples.

### Cortico-cortical and hippocampo-cortical co-ripples couple to thalamo-cortical co-spindles and upstates

Because the data presented above do not support direct driving of cortical ripple co-occurrence by thalamic or hippocampal ripples, we hypothesized that they may be organized by interactions of slower waves, onto which ripples occur. We found that cortical ripples tend to occur at the times of thalamic spindles (Fig.4F,G; see Supp. Fig.13 for cortical activity relative to thalamic ripples). The probability of a cortico-cortical co-ripple occurring during a cortical spindle (on either channel) was significantly increased by 53% when there was a co-occurring anterior thalamic spindle (on any channel) (Fig.4H; *p*=3×10^−69^, *t*(2915)=18.0, one-sided paired *t*-test). Likewise, the probability of a cortico-cortical co-ripple preceding a cortical upstate peak (within 500ms) was significantly increased by 68% when there was a preceding thalamic upstate peak (within 500ms from the cortical upstate peak) (Fig.4H; *p*=3×10^−105^, *t*(2948)=22.7). Thus, these data indicate that projecting thalamic spindles and upstates (but not ripples) help to coordinate cortico-cortical co-ripples (Fig.4F). Similarly, there was a 77% increase in cortico-cortical co-rippling during cortical spindles that co-occurred with a posterior thalamic spindle (Fig.4I; *p*=4×10^−6^, *t*(17)=6.2). However, there was no corresponding significant increase of cortical co-rippling associated with posterior thalamic upstates (*p*=0.87, *t*(27)=1.1).

In performing these same analyses except with respect to hippocampo-cortical co-ripples, the probability of a hippocampo-cortical co-ripple was significantly increased by 31% during anterior thalamo-cortical versus isolated cortical spindles (Fig.4J; *p*=3×10^−5^, *t*(543)=4.0), and significantly by 11% preceding a cortical upstate that was preceded by an anterior thalamic upstate (Fig.4J; *p*=0.003, *t*(544)=2.7).

### Cortico-cortical co-rippling is enhanced during phase-locked thalamo-cortical co-spindles

We showed that if a cortical site spindles with the thalamus during a ripple, it is more likely to co-ripple with another cortical site. We also found that thalamic spindles phase-lock with cortical spindles, consistent with a thalamo-cortical projection of spindles. We therefore hypothesized that cortico-cortical co-rippling is driven by phase-locked thalamo-cortical spindles. We tested if cortico-cortical co-ripple probability is greater between cortical sites that have 10-16Hz spindles phase-locked with the thalamus versus those that do not. We found an increased cortico-cortical co-ripple probability when at least one cortical site had phase-locked spindles with the thalamus (Fig.4J; *p*=4×10^−4^, *t*(1936)=3.3). By contrast, there was no significant increase in hippocampo-cortical co-ripple probability among thalamo-hippocampal (*p*=0.86, *t*(772)=1.1), thalamo-cortical (*p*=0.83, *t*(662)=0.97), or both channel pairs (*p*=0.73, *t*(360)=0.61) that had phase-locked co-spindles versus those that did not. Thus, our data indicate that cortico-cortical co-rippling is further enhanced when thalamo-cortical spindles phase-lock (Fig.4K).

## Discussion

Previous work has shown that human cortical ripples often co-occur and even phase-lock at long separations, posing the question of whether they are synchronized by a subcortical input (Dickey et al., 2022b). The present study investigates if ripples also occur in the human thalamus, and if so whether they, or other thalamic waves, have a role in synchronizing cortical ripples. Indeed, we found that ripples occur in the human anterior and posterior thalamus during NREM, with similar oscillation frequency, occurrence density, and duration as neocortical and hippocampal ripples. The thalamic ripple oscillation frequency was ∼90Hz, similar to human cortical (Dickey et al., 2022a; Dickey et al., 2022b; van Schalkwijk et al., 2022) and hippocampal ripples (Jiang et al., 2019a, b), but lower than the 140-160Hz ripples in rodents (Buzsáki, 2015). We found that, like cortical ripples (Dickey et al., 2022a), thalamic ripples tended to occur after the downstate, and just before the subsequent upstate peak, at the onset of the local sleep spindle.

The term “ripple” has usually been associated with sharpwaves, hippocampus, NREM, and hippocampus-dependent consolidation. However, cortical and hippocampal ripples during both NREM and wakefulness closely share several characteristics, as previously documented (Dickey et al., 2022a). The terms “high-gamma” and “high-frequency oscillation” are too generic to capture the stereotyped phenomenon that we report here since they lack specificity for frequency and duration, do not necessarily refer to discrete events, and may not require distinct oscillations or consistent focal frequencies. These characteristics may be crucial for the generation and function of co-ripples. Thus, the term “ripple” aptly applies to the thalamic events here, and we generally advocate for broadening the term ripple to include events in other sites and states, and with a broader function. Sharpwave-ripples recorded in humans are likely homologous to those in rodents (Jiang et al., 2019a, b; Jiang et al., 2019c). However, human hippocampal ripples during NREM may also occur during spindles. In the current study, as in most other studies of human hippocampal ripples, we did not restrict hippocampal ripples to those associated with sharpwaves, in order to maximize the detection of co-ripples with the thalamus or cortex.

While it is impossible to definitively exclude that our results are not influenced by epilepsy, we carefully sub-selected patients, channels, epochs, and events using iteratively developed manual and automatic methods, in order to exclude epileptic activity. Furthermore, the ripples we detected have the same fundamental characteristics of those detected in microelectrode recordings from the cortex of patients with tetraplegia but not epilepsy (Rubin et al. 2022; Verzhbinsky et al., unpublished) as well as those with sub-cortical tumors but not epilepsy (Cleary et al., unpublished).

Thalamic gamma oscillations, usually at 40Hz but extending to 80Hz, have been recorded in cats and rodents, where they are generated by voltage-gated dendritic Ca^2+^ currents (Pedroarena and Llinás, 1997). This rhythmic firing led to the proposal that the thalamus drives synchronous gamma oscillations across widespread neocortical areas via the widely distributed thalamo-cortical projections of the matrix system (Barth and MacDonald, 1996; Pedroarena and Llinás, 1997; Jones, 2009), but direct evidence is absent. Gamma oscillations have also been reported in mouse and primate visual cortex (Berens et al., 2008). However, these are also lower frequency events (∼50Hz), which appear to have a more restricted anatomical distribution and only synchronize over short distances (Roelfsema, 2023), and may therefore represent a distinct phenomenon.

A direct driving of cortical ripples by thalamic ripples would imply strong synchrony between them. However, thalamic ripples only weakly co-occurred and rarely phase-locked with cortical ripples. These findings, in stark contrast to the robust ripple co-occurrence between cortical sites in this and our previous study (Dickey et al., 2022b), suggest that thalamic ripples do not directly drive widespread forebrain ripple synchrony. Although cortical ripples did not synchronize with thalamic ripples, cortico-cortical co-rippling was strongly promoted by the co-occurrence of spindles or upstates in the thalamus and cortex. Critically, when spindles or upstates coincided between the thalamus and a cortical site, the cortical site was ∼40-50% more likely to co-ripple with another cortical site. Furthermore, spindles preferentially phase-locked between anterior thalamus and anterior cortex, and between posterior thalamus and posterior cortex, as shown in Mak-McCully et al. (2017), which is consistent with anatomical projections from the thalamus. We found that thalamo-cortical spindle phase-locking further enhanced cortico-cortical co-rippling, but neither thalamo-hippocampal nor thalamo-cortical spindle phase-locking enhanced hippocampo-cortical co-rippling. Thus, the thalamus appears to coordinate cortical co-rippling, not by projecting ripples, but by projecting synchronous spindles and upstates. This is consistent with the functional coupling between thalamic and cortical spindles and down-to-upstates in humans (Mak-McCully et al., 2017; Schreiner et al., 2022) and animals (Contreras et al., 1996; Crunelli et al., 2018).

Thalamic spindles and upstates may trigger cortical ripples by depolarizing cortical neurons for ∼50-100ms. Evidence for such depolarizations during NREM has been found in the prominent current sinks and large increases in pyramidal and interneuron firing in microelectrode array recordings from human cortex during upstates and spindles (Cash et al., 2009; Csercsa et al., 2010; Hagler et al., 2018; Dickey et al., 2022a). Rodent hippocampal ripples can be induced by such broad ∼60ms duration depolarizing pulses, via a mechanism involving pyramidal-basket cell feedback loops (“PING”) synchronized by basket–basket cell co-firing (“ING”) (Stark et al., 2014). Further evidence that this mechanism underlies human cortical ripples during NREM is that they are accompanied by strongly phase-locked unit firing, with pyramidal cells leading interneurons (Dickey et al., 2022a).

Previous studies in humans suggest that hippocampal ripples facilitate information transfer to the cortex during sleep (Helfrich et al., 2019; Jiang et al., 2019a, b). Both anterior and posterior thalamic spindles co-occur with cortical spindles (Mak-McCully et al., 2017; Bernhard et al., 2022). We examined the three-way relationships among thalamic, hippocampal, and cortical ripples. And found that thalamic ripples only rarely and weakly co-occur and do not phase-lock with hippocampal ripples. We also confirmed our previous finding that cortical ripples have limited co-occurrences and do not phase-lock with hippocampal ripples (Dickey et al., 2022b). Thus, hippocampal ripples do not appear to be driven by thalamic ripples. However, since hippocampal ripples coordinate with cortical spindles, downstates, and upstates (Jiang et al., 2019a, b), the thalamus could coordinate ripples in the hippocampus and cortex, via a depolarizing modulation as outlined above, rather than via high-frequency oscillatory driving. Indeed, thalamic upstates that project to the cortex or hippocampus were associated with an increased probability of hippocampo-cortical co-ripples.

The thalamus is comprised of many nuclei subserving different physiological functions (Bernhard et al., 2022; Schreiner et al., 2022). The present study distinguishes anterior and posterior thalamus, but further granularity was not possible in these rare recordings with limited sampling and relatively wide spacing between relatively large recording contacts. Therefore, our results only portray general relationships between the anterior and posterior thalamus with the cortex.

Overall, our findings are consistent with cortico-cortical ripple phase-locking arising from direct interactions between cortical neurons. Cortico-cortical ripple synchrony has been proposed to support the binding of specific information encoded in the interacting co-rippling locations (Gray et al., 1989; Dickey et al., 2022b). In such models, not only are oscillations modulated by the interaction of cortical areas through phase-selection (Fries, 2009) and coincidence detection (Salinas and Sejnowski, 2001), but in addition these interactions themselves reinforce the oscillations through re-entrance (Bazhenov et al., 2008; Vicente et al., 2008). A mutually-reinforcing mechanism is consistent with our previous finding of a strong dependence of phase-locking on the co-rippling of several cortical locations simultaneously, beyond the two whose ripple phase-locking is being measured (Dickey et al., 2022b). Because the integration of specific information during binding requires high bandwidth, it may be most effectively transmitted by direct cortico-cortical connections in humans, given that neuroanatomical calculations imply that >95% of synapses on cortical cells are from the cortex (reviewed in Rosen and Halgren (2022)). In particular, the ratio of cortical neurons to thalamo-cortical projection cells is ∼1400:1, based on human thalamic cell counts in Xuereb et al. (1991), cortical cell counts in Azevedo et al. (2009), and the proportion of thalamo-cortical cells in Arcelli et al. (1997). Thus, phase-locking arising from cortico-cortical interactions is consistent with the proposed role of co-ripples in supporting cross-cortical information binding. Our finding that thalamo-cortical spindles and upstates, but not thalamic ripples, enhance cortico-cortical and hippocampo-cortical co-rippling is consistent with the thalamus having a slower, modulatory rather than a high-frequency, direct driving role in promoting co-rippling.

In sum, we provide the first evidence that ripples are generated in the human anterior and posterior thalamus with similar characteristics as hippocampal and cortical ripples. Due to infrequent and weak co-occurrence and phase-locking between ripples in the thalamus and those in the cortex or hippocampus, it is unlikely that thalamic ripples directly drive cortico-cortical or hippocampal-cortical ripple co-occurrence or phase-locking. Rather, we found evidence that thalamic upstates, spindles, and especially phase-locked spindles coordinate the co-occurrence of ripples in the cortex. This may help to organize replay that underlies the consolidation of memories across widespread areas in the cortex.

## Methods

### Patient selection, intracranial recordings, and data pre-processing

Data from 13 patients (8 female, 39.5±11.5 years old) with pharmaco-resistant focal epilepsy undergoing intracranial recording for approximately one to two weeks to localize seizure onset zones prior to surgical resection were included in this study (Supp. Table 1). The patients included in this study underwent extensive clinical non-invasive evaluation that indicated that they had medication-resistant focal epilepsy that could be treated surgically. However, it is difficult to determine the location of the epileptic foci non-invasively, and these foci vary across patients. Therefore, electrode targeting was performed in a patient-specific manner in order to target each patient’s suspected epileptic foci.

As reviewed by (Gadot et al., 2022), implantation of the thalamus together with cortical sites is a clinically-accepted procedure in the pre-surgical evaluation of partial epilepsy. Partial epilepsy is typically a network phenomenon, and the thalamus is the most common extra-temporal site that is recruited by temporal lobe seizures (Rosenberg et al., 2006; Pizzo et al., 2021; Yan et al., 2022). Thus, thalamic recordings can provide important information regarding the sequence of engagement of different cortical areas during the evolution of the seizure discharge. Although the thalamus is not a target for resection, it can be a target for FDA-approved deep brain stimulation therapy. Through an IRB-approved process, we have prospectively recruited adults (>18 years old) undergoing SEEG investigation for the localization of seizure foci to participate in the thalamic electrophysiological recordings. Fully written informed consent was obtained from all patients before the thalamic SEEG implantation. Implantation and consent procedures for the patients with anterior thalamic electrodes have been described in detail in our prior study (Chaitanya et al., 2020). Those procedures for the posterior thalamic electrodes were highly similar. In summary, one of the clinically indicated depth electrodes sampling the insula operculum was advanced deeper to target thalamic nuclei. Thus, we avoided using additional depth electrodes for exclusive research implantation and mitigated the risk of bleeding, which is known to be higher with an increased number of depth electrodes implanted. None of the patients had a thalamic bleed. All patients in the study gave fully informed written consent for their data to be used for research. This study was approved by the local Institutional Review Boards at the University of Alabama, Birmingham (patients 1-10 with anterior thalamic electrodes) and the French Institute of Health (patients 11-13 with posterior thalamic electrodes).

Patients were only included in this study if they had no prior brain surgery and normal background LFPs except for infrequent epileptiform activity. A total of 15 patients with thalamic electrodes were rejected due to abnormal LFPs if they met any of the following criteria: 1) more than one automatically detected (see below) putative inter-ictal spike per minute on average; 2) more than one >1 mV artifact per minute on average as determined visually across ≥60 minutes of data. Patient 1-10 recordings were collected from PMT electrodes (PMT Corporation, Chanhassen, Minnesota, USA) with a Natus Quantum amplifier (Natus Medical Incorporated, Pleasanton, CA, USA) at 2048Hz sampling with a 0.016-683Hz bandpass. Patient 11 recordings were collected at 512Hz with a 160Hz bandpass. Patient 12-13 recordings from DIXI electrodes (DIXI Medical, Marchaux-Chaudefontaine, France) were collected at 1024Hz with a 0.16-340Hz bandpass. PMT electrodes were 0.8mm diameter with 10-16 2mm long contacts separated by 1.5-3 mm (∼150 contacts/patient). Data preprocessing was performed in Matlab 2019b (MathWorks, Natick, MA, USA) and LFPs were inspected visually using the Fieldtrip toolbox (Oostenveld et al., 2011). Data sampled above 1000Hz were downsampled to 1000Hz with anti-aliasing at 500Hz. All data were notch filtered at 60Hz with 60Hz harmonics (patients 1-10) or 50Hz with 50Hz harmonics (patients 11-13) up to the Nyquist frequency.

### Channel selection

Channels were bipolar derivations of adjacent contacts to ensure measurement of LFPs. Channels were only included if their contacts were in non-epileptogenic, non-lesioned thalamus as well as cortex, and in 9 of the 13 patients also hippocampus. Furthermore, cortical and hippocampal channels were excluded if they shared a common contact with another bipolar channel that was included in the analysis, whereas all thalamic bipolars that met criteria were included in order to maximize sampling from the many nuclei in the thalamus. In order to ensure that recordings reflect local activity rather than that volume conducted from elsewhere, we restricted recordings to the gray matter. Initial selection of bipolar contacts that may be in the gray matter was made by using the co-registered pre-operative MRI and post-operative CT to localize contacts. However, the cortical ribbon is thin (∼3 mm in humans), so in addition to using the co-registered pre-operative MRI and post-operative CT to localize contacts, we used physiological measurements to select among possible disjoint bipolar pairs those with the largest spontaneous LFP amplitude. Specifically, channels were manually selected based on NREM averages of narrowband delta (0.5-2Hz) and high-gamma (70-190Hz) analytic amplitudes. Cortical channels selected had an average delta peak amplitude greater than 40µV, hippocampal channels had a delta amplitude greater than 15µV, and thalamic channels had a delta amplitude greater than 5µV. In addition, they had significant correlations (*p*<0.05) between the low-pass delta waveform and the high gamma (70-190Hz) analytic amplitude. Delta was used because its power is increased during non-rapid eye movement sleep that is characterized by slower frequency oscillations including the slow oscillation and K-complexes. High gamma was used because its power reflects local neuron spiking generated by cells in the gray matter. Both are present in virtually all cortical and hippocampal bipolar channels during NREM sleep stage N3. Channels were then visually inspected during NREM periods to ensure they did not have frequent interictal spikes or artifacts and had a normal appearing broadband signal. Among 2145 bipolar channels (1157 left sided) across the patients in the study, a total of 336 channels (110 left sided) were included in the study. Among the 39 thalamic bipolar channels across the patients in the study, 30 channels were included in the study because they had normal appearing LFPs without evidence of frequent inter-ictal spiking. Note that the majority of channels were excluded because they were not in thalamic, cortical, or hippocampal gray matter, and thus would not record meaningful local neuronal activity, or in order to keep bipolar derivations disjoint.

### Channel localization

Cortical surfaces were reconstructed using the pre-operative T1-weighted structural MR volume with the FreeSurfer recon-all pipeline (Fischl, 2012). Atlas-based automated parcellation (Fischl et al., 2004) was utilized to assign anatomical labels to cortical surface regions according to the Destrieux et al. (2010) atlas, and subsequently amalgamated into cortical parcels per Desikan et al. (2006). For analyses involving anterior versus posterior cortex, anterior cortex included Desikan parcels that comprise the orbitofrontal cortex (frontal pole, lateral orbitofrontal, medial orbitofrontal, pars orbitalis), prefrontal cortex (caudal middle frontal, pars opercularis, pars triangularis, rostral middle frontal, superior frontal), and cingulate cortex (caudal anterior cingulate, rostral anterior cingulate, posterior cingulate). Posterior cortex was comprised of the remaining cortical parcels.

To localize the SEEG contacts, the post-implant CT volume was co-registered to the pre-implant MR volume in standardized 1mm isotropic FreeSurfer space with the general registration module (Johnson et al., 2007) in 3D Slicer (Fedorov et al., 2012). Next, each contact’s position in FreeSurfer coordinates was determined by manually marking the contact centroid as visualized in the co-registered CT volume. Each transcortical bipolar channel was assigned an anatomical parcel from the above atlas through determining the parcel identities of the white-gray surface vertex nearest to the midpoint of the adjacent contacts. Automated segmentation assigned a nucleus to each voxel of the MR volume in the thalamus (Iglesias et al., 2018), which was confirmed by visual comparison with the Allen Brain Atlas (https://atlas.brain-map.org/). The 24 thalamic bipolar channels included the following nuclei (in descending frequency): AnteroVentral, Ventral Anterior, Reticular, Ventral Lateral, Fasciculus, MedioDorsal, and CentroMedian (Supp. Table 1). Due to possible inaccuracies in registration and limitations of visible nuclear boundaries, assignment of contacts to particular nuclei should be considered approximate. Transcortical bipolar locations were registered to the fsaverage template brain for visualization through spherical morphing (Fischl et al., 1999). All contacts marked as thalamic or hippocampal were confirmed to localize to these respective structures through visual inspection of the co-registered pre-operative MRI and post-operative CT. The posterior limit of the uncal head was used as a boundary to define anterior vs. posterior hippocampus (Poppenk et al., 2013; Ding and Van Hoesen, 2015).

### Sleep staging and epoch selection

Sleep staging was performed to select NREM stages N2 and N3 consistent with Silber et al. (2007). Data were bandpassed at 0.5-2Hz and then Hilbert transformed to obtain the narrowband delta analytic amplitudes. NREM epochs were selected for analysis when the cortical channels had increased delta analytic amplitude during overnight hours (8:00 PM-8:00 AM). These epochs subsequently underwent visual confirmation that they contained frequent and prominent slow wave oscillations and spindles characteristic of NREM stages N2 and N3. Epochs were only retained when they were absent of frequent inter-ictal spikes or artifacts. Notably, disruptions to sleep (e.g., for vitals assessment) may occur while patients are on the epilepsy monitoring unit despite our methods being designed to select for the most normal overnight NREM sleep in these patients.

### Time-frequency analyses

Time-frequency plots of ripple-triggered spectral power were generated from the broadband LFP using EEGLAB (Delorme and Makeig, 2004). Power was calculated from 1-500Hz by averaging the fast Fourier transforms with Hanning window tapering locked to the centers of 5000 randomly sub-selected ripples per channel. Each 1Hz frequency bin was normalized to the mean power at −2 to −1.5s and then masked with two-tailed bootstrapped significance (*N*=200 shuffles) with *p*-values FDR-corrected with α=0.05 using a −2 to −1.5s baseline. Grand average time-frequency plots were produced by averaging across the channel average time-frequency plots.

### Ripple detection

Ripples were detected in the thalamus, cortex, and hippocampus based on previously published methods (Dickey et al., 2022a; Dickey et al., 2022b). For each channel, data were bandpassed at 70-100Hz with a 6^th^ order forward-reverse Butterworth filter (zero-phase shift) and peaks in the analytic amplitude of the Hilbert transform were selected if they were at least 3 standard deviations above the channel mean. Event onsets and offsets were found on both sides of the peak when the 70-100Hz analytic amplitude decreased below a z-score of 0.75. Events were retained if they had at least 3 oscillation peaks in the 120Hz lowpass, determined by moving a 40ms window in 5ms increments ±50ms relative to the midpoint of the ripple, requiring at least 1 window to have at least 3 peaks. Adjacent events within 25ms were merged and ripple centers were determined as the time of the largest positive peak in the 70-100Hz bandpass.

To exclude epileptiform activity and artifacts, events were rejected when the absolute value of the z-score of the 100Hz highpass was greater than 7. Events were also rejected if they were within ±500ms from putative interictal spikes (detection criteria described below). Furthermore, events were rejected if they coincided with a putative interictal spike detected on any channel to exclude events that could be coupled across channels due to epileptiform activity. Lastly, events were rejected if the largest peak-to-valley or valley-to-peak absolute amplitude in the broadband LFP was 2.5 times greater than the third largest in order to exclude events that had only one prominent cycle or deflection. The average broadband LFP and average time-frequency plots locked to ripple centers as well as multiple individual broadband LFP ripples were visually examined in each channel from each patient to confirm there were multiple distinct cycles in the 70-100Hz band, without contamination by possible artifacts or epileptiform transients. The oscillation frequency of each ripple was computed as follows:

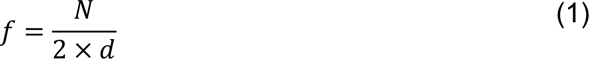

Where *N* is the number of 70-100Hz zero crossings (i.e., half cycles, with fractional cycles determined based on the remaining phase angle over π), and *d* is the duration of the ripple.

### Interictal spike rejection

Ripples were excluded from analysis if they were within 500ms from putative interictal spikes. Putative interictal spikes were detected by finding a high frequency score, which was the 20ms boxcar smoothed 70-190Hz analytic amplitude, and a spike template score, which was computed as the cross-covariance with a template interictal spike. A putative interictal spike was marked when the sum of the high frequency score, weighted by 13, and the spike template score, weighted by 25, exceeded 130. These thresholds were determined through extensive visual examination of many hundreds of events across multiple channels from multiple patients.

### Downstate, upstate, and spindle detection

Downstates and upstates were detected according to previously published methods (Jiang et al., 2019a, b; Dickey et al., 2021; Dickey et al., 2022a). Data were bandpassed from 0.1-4Hz (delta) and the top 10% of amplitude peaks between consecutive zero crossings within 0.25-3s were identified. The polarity of each bipolar signal was inverted if needed so that downstates were negative and upstates were positive. This was done by confirming that the average 70-190Hz analytic amplitude ±100ms (cortex) or ±250ms (thalamus) around peaks was greater for upstates than downstates for each channel. Broadband high gamma (70-190Hz) was used as it is a proxy for multi-unit activity that is enhanced during the upstate and silenced during the downstate. Further determination of thalamic polarities, which may be difficult given the small magnitude of high gamma (Mak-McCully et al., 2017), was performed by visually inspecting the spindle-locked delta signal with respect to the high gamma analytic amplitude. Polarity was determined such that the high gamma envelope increased during the rising phase of the delta signal (i.e., the down-to-upstate transition).

Spindles were also detected according to a previously published method (Hagler et al., 2018; Dickey et al., 2021; Dickey et al., 2022a). Data were bandpassed at 10-16Hz, and the absolute values were smoothed with convolution using a tapered 300ms Tukey window, then median values were subtracted from each channel. Data were normalized by the median absolute deviation, and spindles were identified when peaks exceeded 1 for a minimum of 400ms. Onsets and offsets were marked when these amplitudes fell below 1. Putative spindles coinciding with large increases in lower (4-8Hz) or higher (18-25Hz) power were rejected to exclude broadband events and theta bursts, which may overlap with slower spindle frequencies (Gonzalez et al., 2018).

### Ripple timing and co-occurrence analyses

Timing relationships between ripples and other sleep waves were computed as previously described (Dickey et al., 2022a). Times of ripple centers relative to spindle onsets, downstate peaks, and upstate peaks on the same channel were determined for each channel. Event counts were calculated in 50ms bins ±1500ms around cortical ripple centers. Time histogram plots were smoothed with a 50ms Gaussian window with σ=10ms.

Cortical ripples were considered to co-occur (“co-ripple”) if they overlapped for at least 25ms in two different cortical bipolar channels (no cortical bipolar channels shared a common contact). A cortical co-ripple was determined to co-occur with an isolated cortical spindle if its time center occurred during a spindle on either cortical channel without a thalamic spindle onset preceding the cortical spindle onset within 500ms on any thalamic channel. A cortical co-ripple was determined to co-occur with a thalamo-cortical spindle if there was a thalamic spindle on any thalamic channel with an onset within 500ms preceding the cortical spindle onset. Probabilities were computed as the number of co-ripples divided by the total time of cortical spindling, including only the cortical spindles selected for each condition as described above. This same method was used to compute the probability that a cortical co-ripple occurred within 500ms preceding isolated cortical upstates compared to thalamo-cortical upstates, where the thalamic upstate peak preceded the cortical upstate peak within 500ms.

To compute the probability of cortico-cortical (CX1-CX2) co-rippling given cortical (CX3) rippling, we divided the instantaneous probability of CX1-CX2-CX3 co-rippling by the instantaneous probability of CX3 rippling. Likewise, to compute the probability of CX1-CX2 co-rippling given thalamic (TH) rippling, we divided the instantaneous probability of TH-CX1-CX2 co-rippling by the instantaneous probability of TH rippling. For each CX1-CX2 pair, CX3 and TH was a different, randomly selected channel. Co-rippling required ≥25 ms overlap.

### Phase-locking analyses

Phase-locking of co-occurring ripples/spindles at different sites was evaluated using the phase-locking value (PLV), which is an instantaneous measure of phase consistency that is independent of signal amplitudes (Lachaux et al., 1999). PLV time courses were determined using the analytic angles of the 70-100Hz (for ripples) or 10-16Hz (for spindles) bandpass (6^th^ order zero-phase shift forward-reverse filtered) of each channel pair when there were at least 40 co-ripples that overlapped for at least 25ms or 40 co-spindles that overlapped for at least 200ms. PLVs were computed in 1ms timesteps relative to the temporal centers of the co-ripples/spindles. A null distribution was generated by randomly selecting 500 times within −10 to −2s preceding each co-ripple/spindle center. Pre-FDR *p*-values were computed by comparing the observed vs. null distributions in 5ms bins within ±50ms around the co-ripple centers or within 50ms bins within ±250ms around the co-spindle centers. *P*-values were then FDR-corrected across bins and channel pairs. A channel pair was determined to have significant ripple phase-locking if at least 2 consecutive bins had post-FDR *p*<0.05. PLV plots were Gaussian smoothed with a window of 10ms and σ=2ms.

### White matter controls

Bipolar channels, recorded from the electrodes implanted into the thalamus, which localized to the white matter lateral to the thalamic bipolar channels included in the study were selected for control analyses. One white matter bipolar channel that met these criteria was identified in patients 2, 3, 4, 7, and 10. To test for effects of volume conduction, the mean white matter channel LFP relative to thalamic ripple centers was computed. To test for ripples in the white matter channels, ripple detection was performed with the same criteria as described above using the thalamic channel thresholds.

### Experimental design and statistical analyses

Statistical analyses were performed with *α*=0.05 and all *p*-values involving multiple comparisons were FDR-corrected (Benjamini and Hochberg, 1995) across all channels from all patients unless specified otherwise. Random shuffling statistics were performed with *N*=200 iterations per channel or channel pair. Paired and two-sample *t*-tests were used to compare distribution means. To determine if a peri-event time histogram was significant for a given channel or channel pair, a null distribution was found by shuffling the events relative to the ripples within ±1.5s 200 times per channel pair or channel. *P*-values were computed by a comparison of the observed and null distributions for each bin over ±25ms for ripples or ±50ms (250ms Gaussian smoothed with σ=50ms) for other sleep waves. All channel pairs from all patients were included regardless of the numbers of events (co-occurring or independently) for these pair-wise statistical analyses. These *p*-values were then FDR-corrected across all bins across all channels/channel pairs. A channel/channel pair had a significant modulation if 3 or more consecutive bins had post-FDR *p*<0.05. A one-sided Wilcoxon signed-rank test was used to test for significance of modulation across all channels or channel pairs using the observed versus mean null distribution bin values within −500-0 ms and 0-500 ms or −500-500 ms, respectively. Whether a given channel/channel pair had a leading/lagging relationship between events was determined using a two-sided binomial test to compare the counts in the 250ms preceding vs. following *t*=0 with an expected value of 0.5. Whether a channel pair had significantly greater co-rippling compared to chance was determined by randomly shuffling (*N*=200 iterations) the inter-ripple ripple intervals in moving non-overlapping 5 min windows on each channel and comparing the observed vs. null number of co-occurrences. These *p*-values were FDR-corrected across all channel pairs. Conditional probabilities of cortico-cortical relationships were computed for both P(A|B) and P(B|A). Comparisons of proportions (e.g., proportions of significant channel pairs for various conditions) were performed with a χ^2^ test of proportions. For a given pair-wise analysis, e.g., cortico-cortical ripple co-occurrences, the strength of the relationship was considered to be weak if increased by <25% from chance, or was otherwise considered to be strong. Likewise, the relationship was considered to be infrequent if <25% of channels were significant, or was otherwise considered to be frequent. For analyses of spindle- and upstate-associated changes in co-rippling, channel pairs were excluded from statistical analysis if there were no co-ripples in either condition being compared or if there was less than 15s of total cortical spindling time.

To test for differences of ripple characteristics (i.e., density, amplitude, frequency, duration) between different regions, we used the following linear-mixed effects model formula to control for patient as a random effect, with subsequent FDR-correction for multiple comparisons (across all channels from all patients):

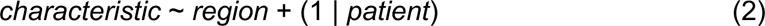

### Data accessibility

The de-identified raw data that support the findings of this study are available from the corresponding authors upon reasonable request provided that the data sharing agreement and patient consent permit that sharing.

### Code accessibility

The code that support the findings of this study are available from the corresponding authors upon reasonable request.

## Author contributions

C.W.D., S.S.C., and E.H. designed the study. P.Y.C. and S.P. collected the data. C.W.D., I.A.V., S.K., B.Q.R., and C.E.G. analyzed the data. C.W.D. and E.H. wrote the manuscript. All authors reviewed and revised the manuscript. E.H. and S.P. supervised the work.

## Acknowledgements

We thank Emília Tóth for assisting with data collection. We also thank Adam Niese, Daniel Cleary, and Jacob Garrett for their support. This work was supported by NIMH (1RF1MH117155-01, T32 MH020002) and ONR-MURI (N00014-16-1-2829).

