## Supplemental Information for "Thalamic spindles and upstates, but not ripples, coordinate cortico-cortical and hippocampo-cortical co-ripples in humans"

**Patient 4**

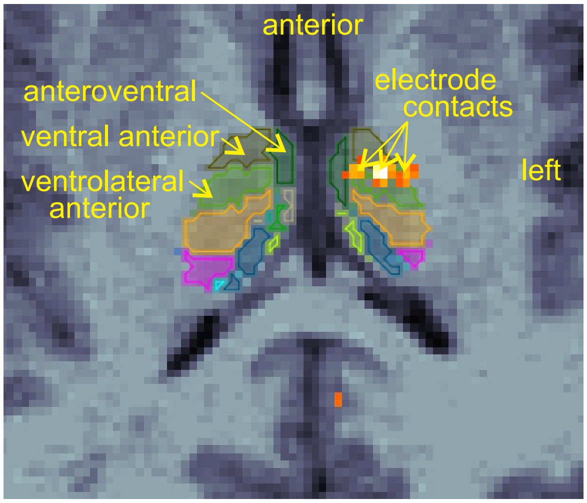


**Supplemental Figure 1. Example thalamic contact localizations**. Pre-operative MR axial section in grayscale with co-registered post-operative CT showing left sided thalamic contacts in orange from a representative patient. Overlaid outlines show thalamic nuclei as estimated by automated segmentation of the T1-weighted MR volume (Iglesias et al., 2018). CT, computed tomography; MR, magnetic resonance. See Supplemental Table 1 for thalamic channel localizations of each patient.

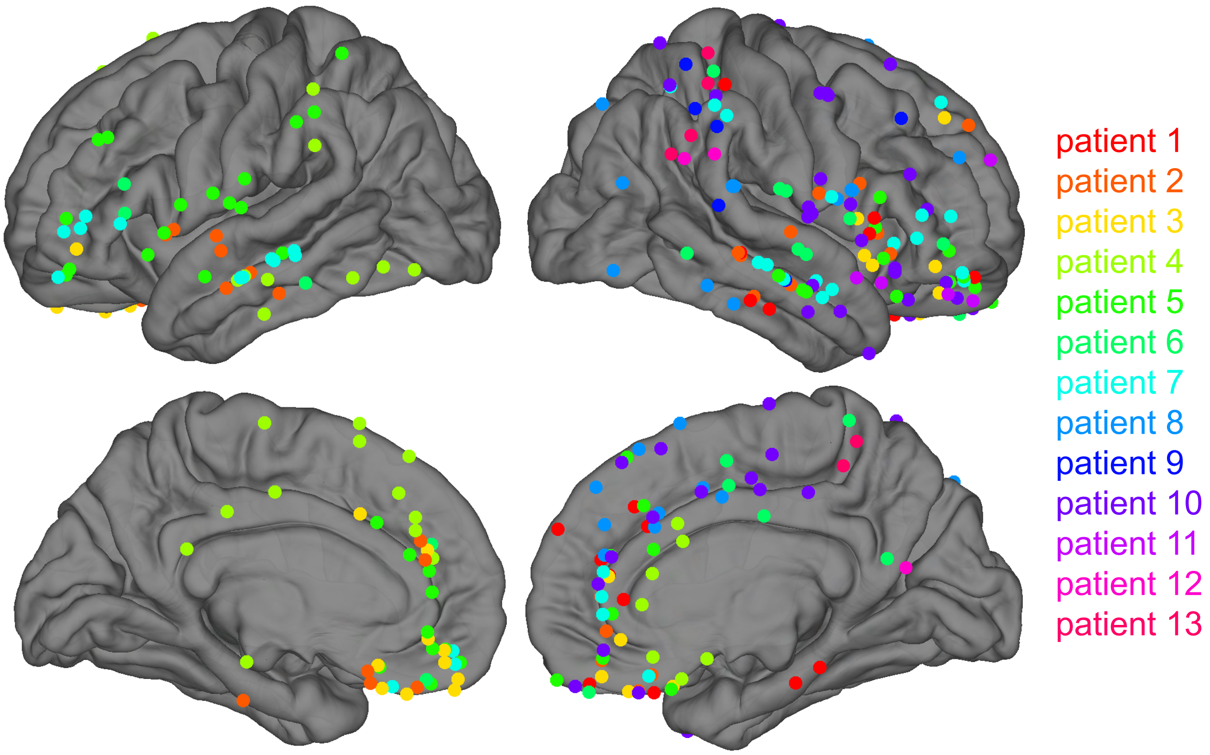


**Supplemental Figure 2. Cortical channel maps.** Markers show centers of all bipolar channels (*N*=275) from all patients (*N*=13). Further anatomical information, including thalamic and hippocampal localizations, is provided in Supplemental Table 1.

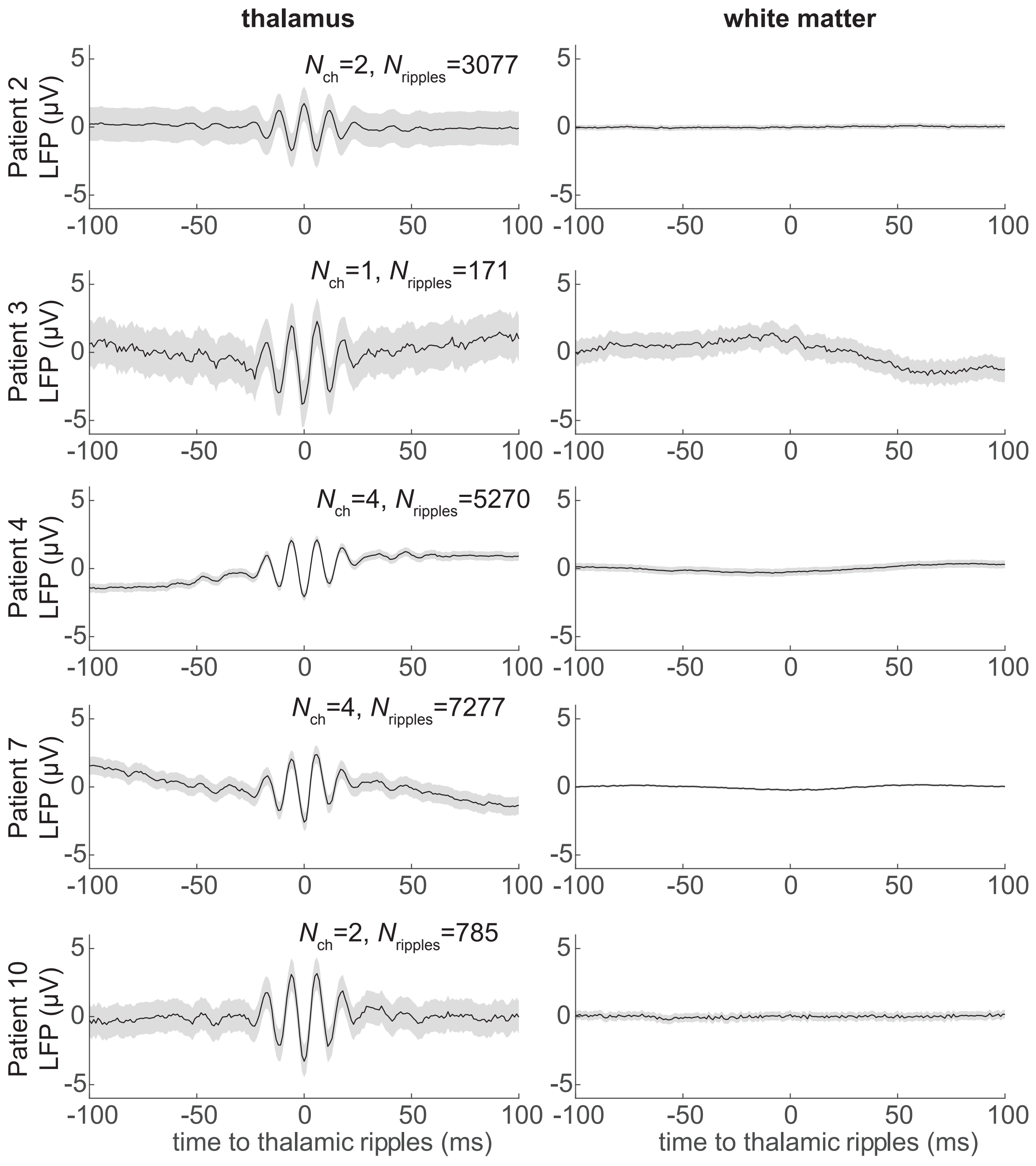


**Supplemental Figure 3. Detection of thalamic ripples is not due to volume conduction.** Average and SEM LFP (left column) and white matter LFP (right column) time-locked to thalamic ripples. All recordings are bipolar referenced in order to ensure focal measurement of LFPs. Note the prominent ripple oscillation that localizes to thalamic gray matter but is not present in the adjacent white matter. LFP, local field potential; SEM, standard error of the mean.

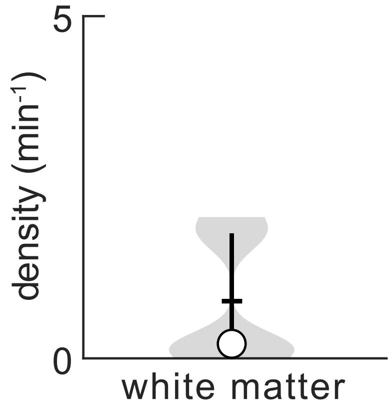


**Supplemental Figure 4. Ripples are almost never detected in white matter adjacent to the thalamus.** Density (frequency of occurrence) of events detected in bipolar channels that localize to the white matter adjacent to the thalamus. Recordings were obtained from the same probes with medial contacts implanted into the thalamus. The ripple densities in the white matter were 4% of those in the cortex as reported in Fig. 1E ($p=5\times10^{-10}$, $t(27)=9.4$; linear mixed-effects with patient as random effect), indicating that the ripples included in this study localize to gray matter and are not due to volume conduction, noise, or artifact. Horizontal lines, means; circles, medians; vertical lines, interquartile ranges.

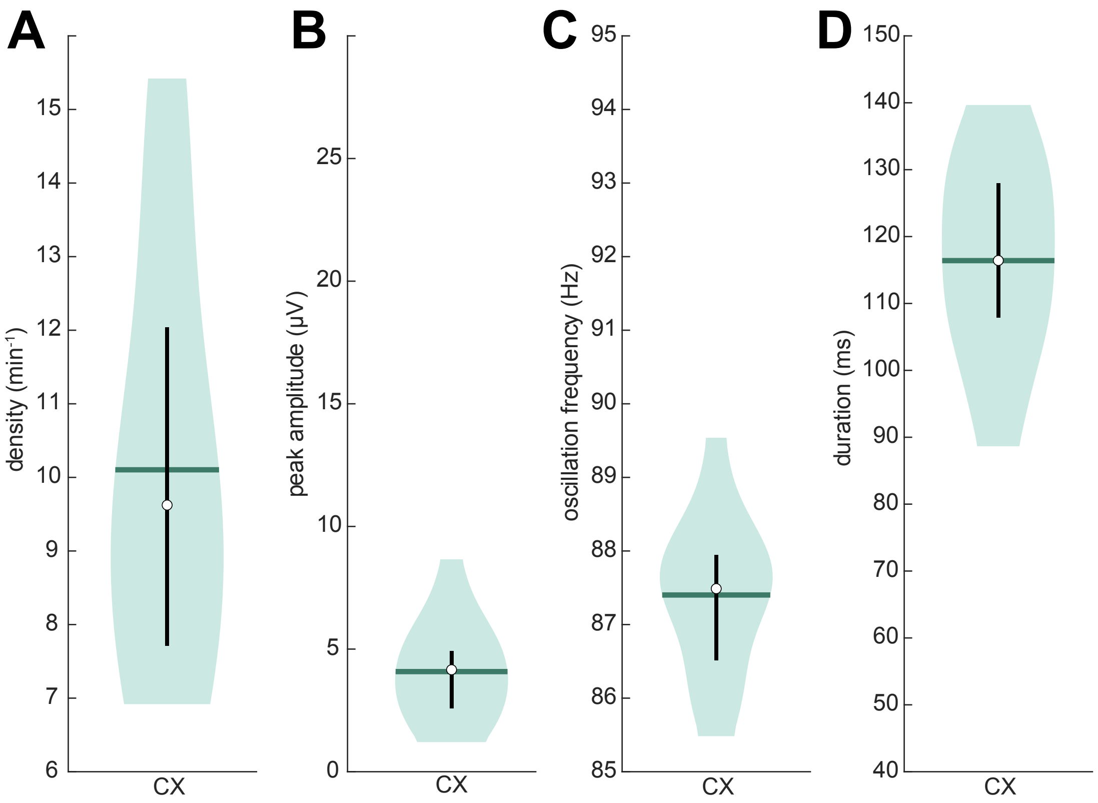


**Supplemental Figure 5. Characteristics of cortical ripples from patients with posterior thalamic recordings.** (**A-D**) Cortical ripple densities (**A**), peak 70-100 Hz analytic amplitudes (**B**), oscillation frequencies (**C**), and durations (**D**) during NREM across all channels (*N*=14 from patients 11-13). Horizontal lines, means; circles, medians; vertical lines, interquartile ranges. CX, cortex.

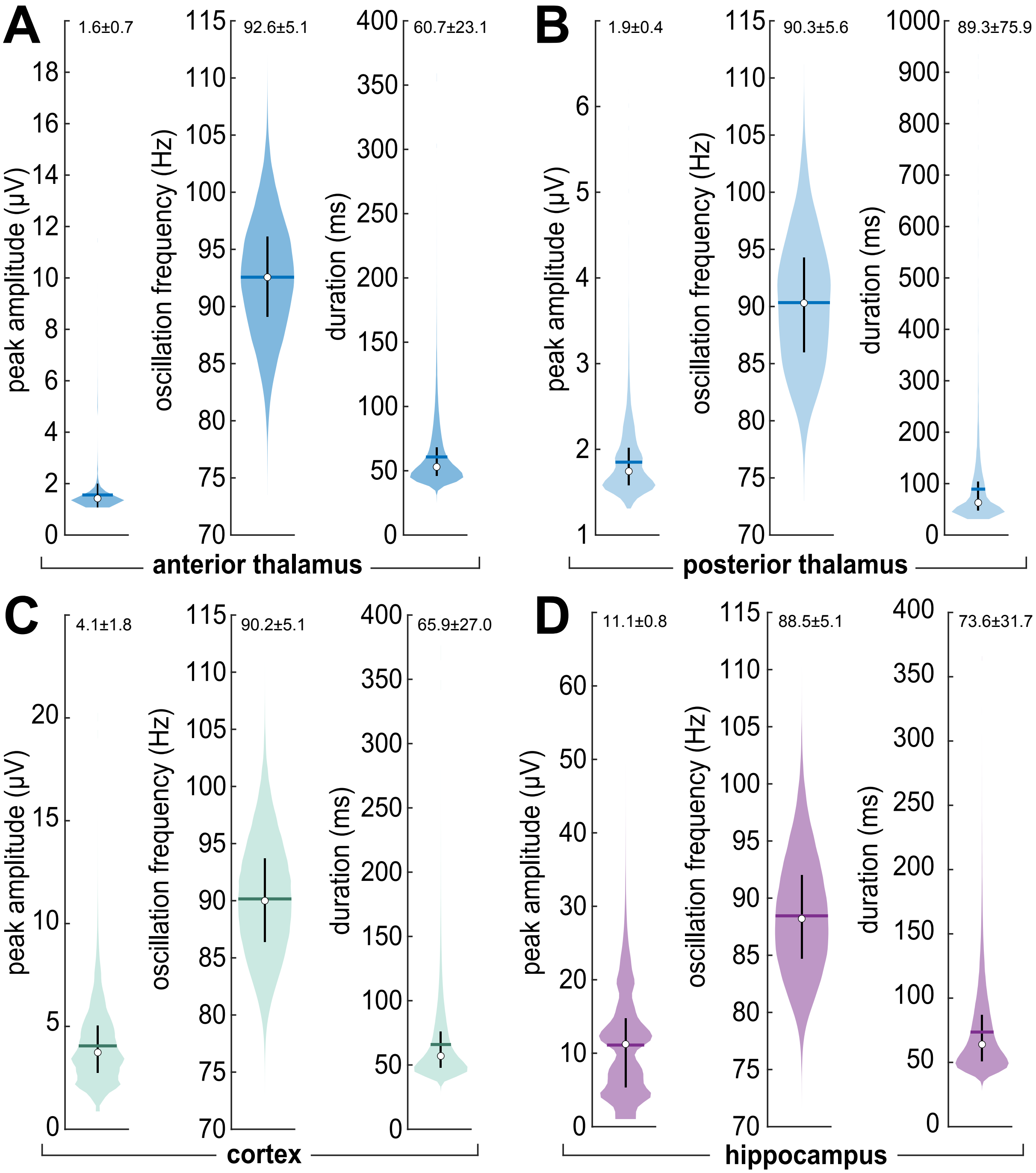


**Supplemental Figure 6. Thalamic, cortical, and hippocampal ripple characteristics across individual ripples. (A-D)** Anterior thalamic (**A**), posterior thalamic (**B**), cortical (**C**), and hippocampal (**D**) peak 70-100 Hz analytic amplitudes, oscillation frequencies, and durations across all ripples ($N_{aTH}$=88,320, $N_{pTH}$= 8,996, $N_{CX}$= 857,420, $N_{HC}$= 135,646) during NREM. Values above each plot are mean and standard deviation. Horizontal lines, means; circles, medians; vertical lines, interquartile ranges. aTH, anterior thalamus; CX, cortex; HC, hippocampus; pTH, posterior thalamus.

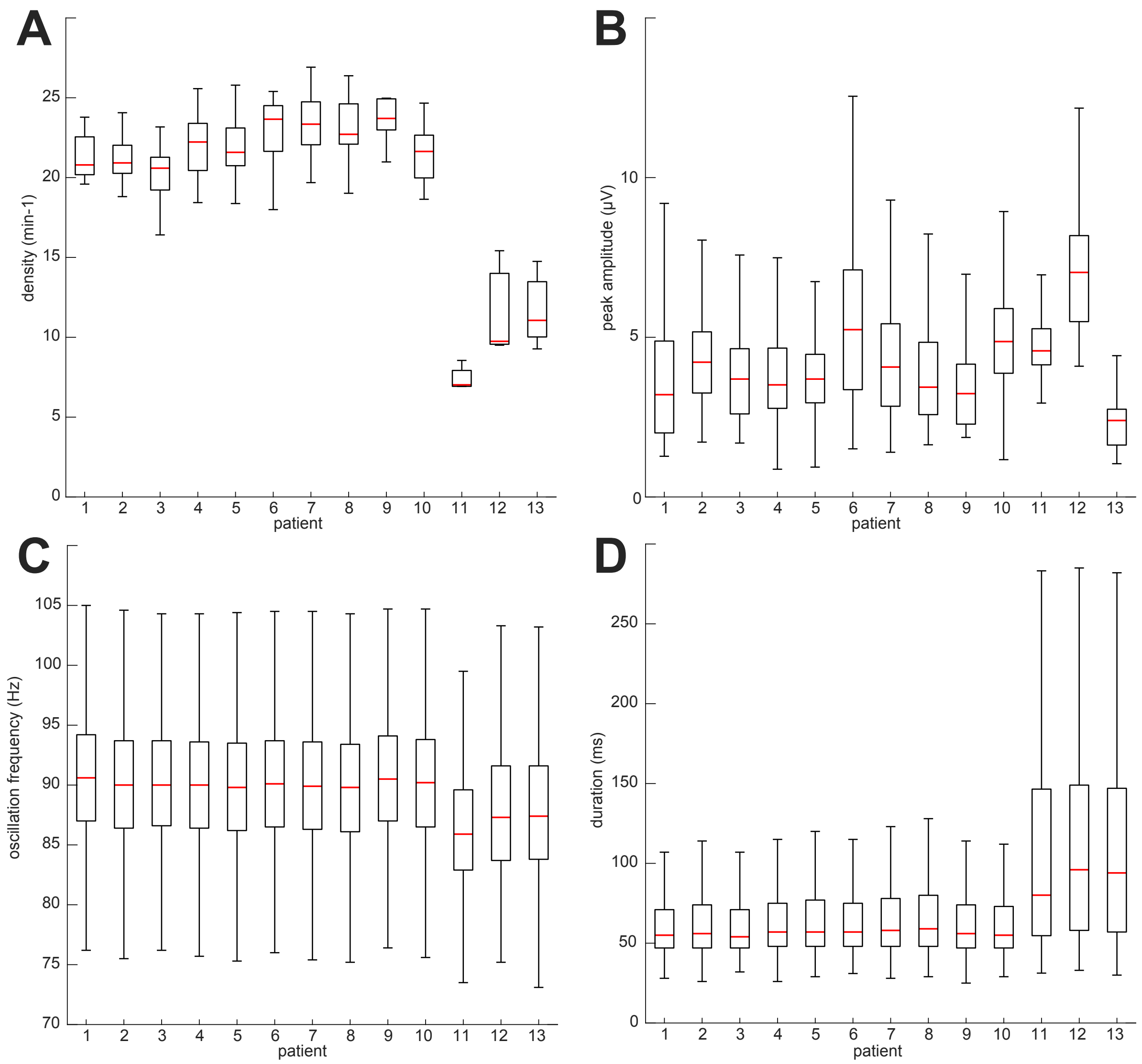


**Supplemental Figure 7. Thalamic ripple characteristics by patient. (A-D)** Thalamic ripple densities (**A**), peak 70-100 Hz analytic amplitudes (**B**), oscillation frequencies (**C**), and durations (**D**) across ripples from each patient. Patients 1-10 are anterior thalamus and 11-13 are posterior thalamus. Boxes show interquartile ranges, horizontal lines indicate medians, and whiskers represent 1.5 ×interquartile range.

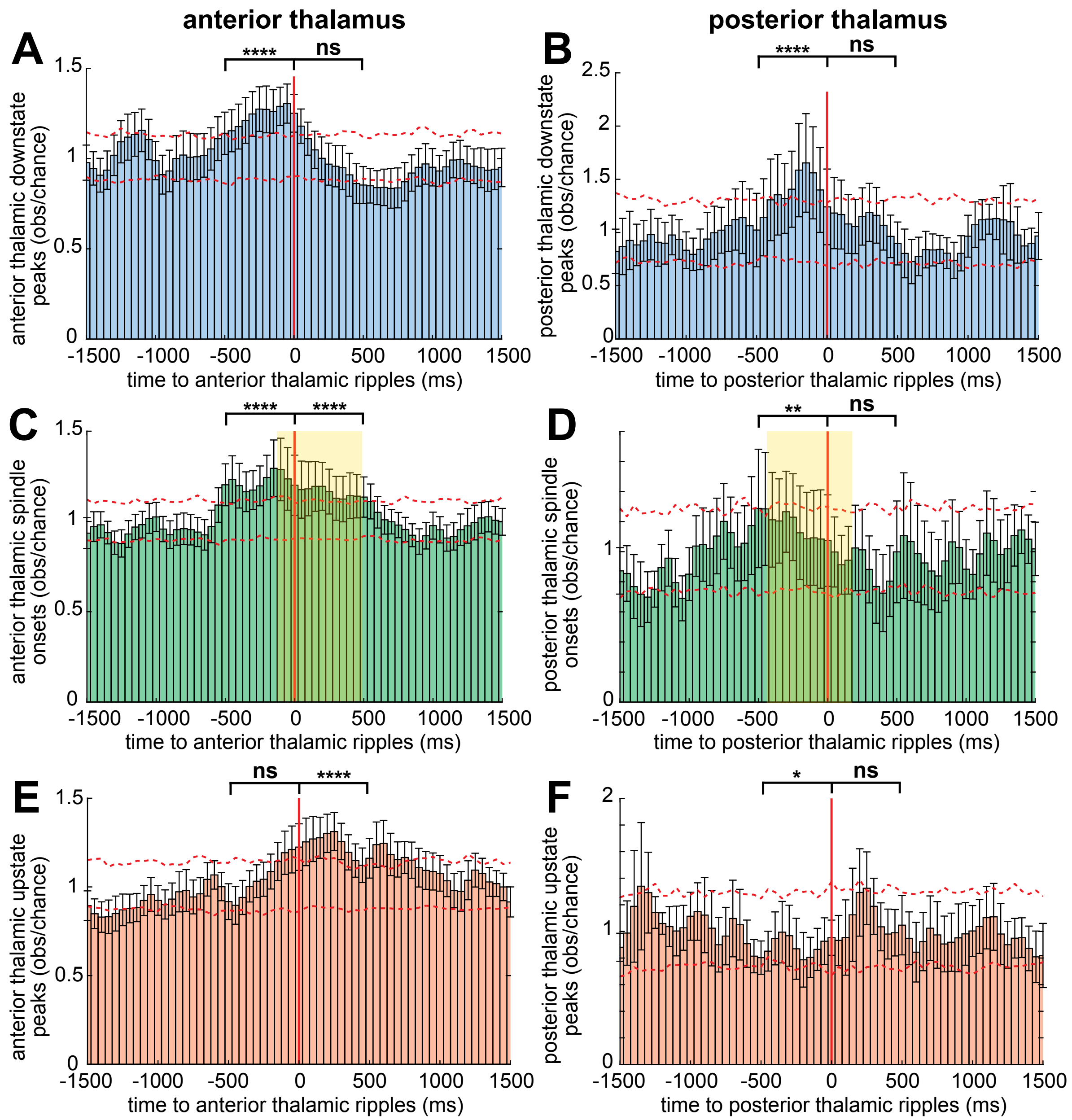


**Supplemental Figure 8. Thalamic ripples occur on the local down-to-upstate transition during spindles.** (**A**) Average and SEM times of anterior thalamic downstate peaks relative to anterior thalamic ripples. (**B**) Same as A except posterior thalamus. (**C-D**) Same as A-B except spindle onsets. Shaded boxes denote average spindle interval. (**E-F**) Same as A-B except upstate peaks. Data are from all channels from all patients. Channels exclusively with significant modulations are depicted in Fig. 1. Dashed error shows 99% confidence intervals of the null distribution. *P*-values computed using a Wilcoxon ranked-sum test to compare the modulation amplitude within -500 to 0 ms and 0 to 500 ms across bins and channels for observed values versus null mean values. ns=non-significant, **p*<0.05, ***p*<0.01, ****p*<0.001, *****p*<0.0001.

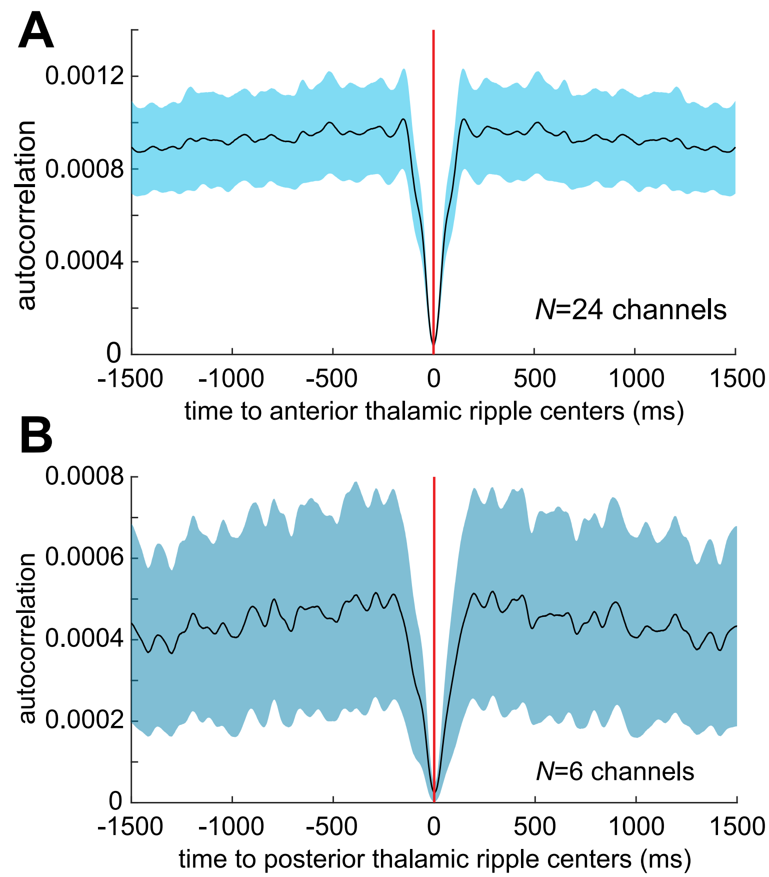


**Supplemental Figure 9. Thalamic ripple autocorrelation.** (**A-B**) Average and SEM within-channel autocorrelation of anterior (**A**; *N*=24 channels) and posterior (**B**; *N*=6 channels) thalamic ripples.

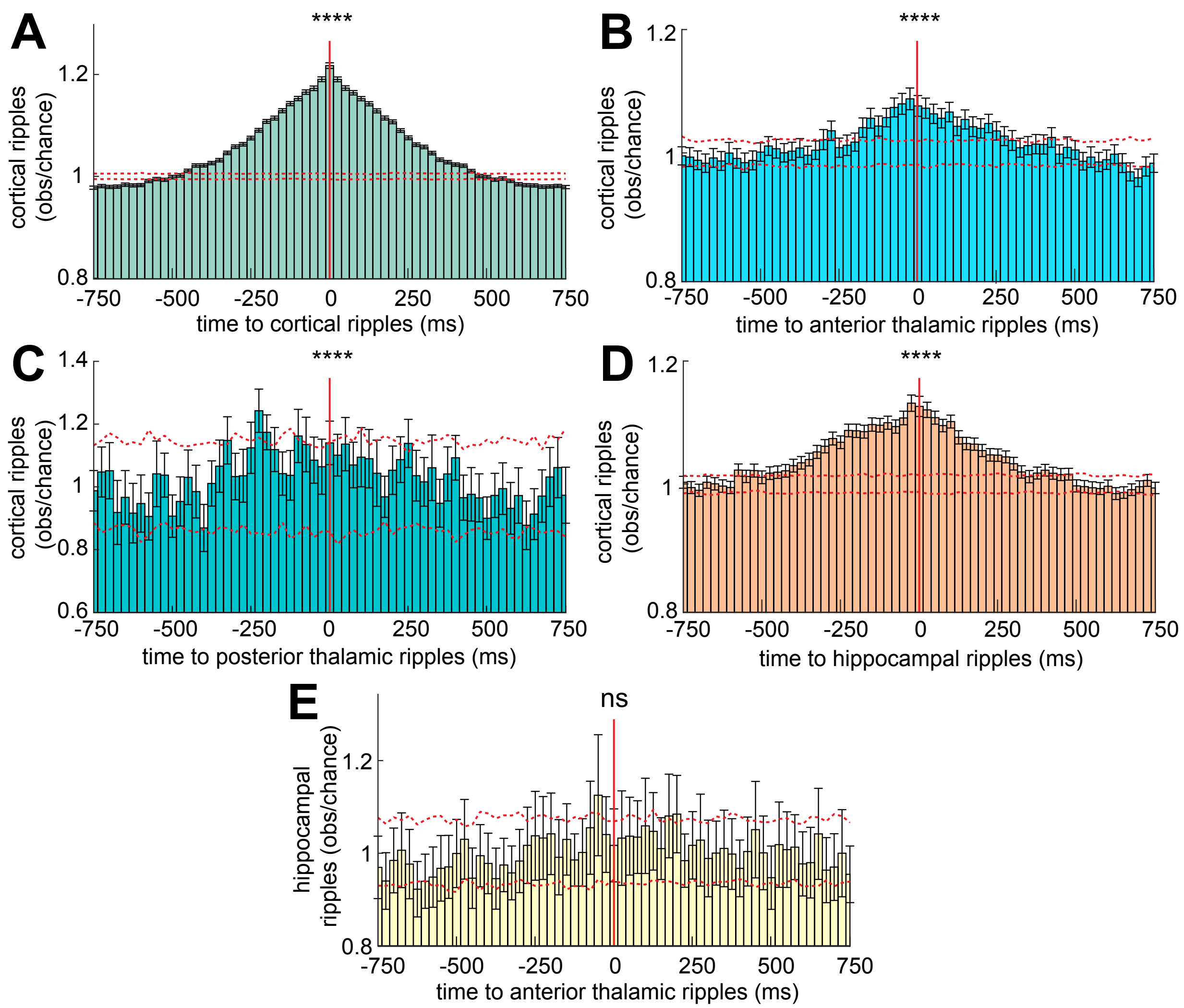


**Supplemental Figure 10. Thalamic ripples infrequently and weakly co-occur with cortical and hippocampal ripples during NREM.** (**A**) Cortical ripples on one channel relative to those on another (*N*=7796 channel pairs). (**B**) Cortical relative to anterior thalamic ripples (*N*=649). (**C**) Cortical relative to posterior thalamic ripples (*N*=26 channel pairs). (**D**) Cortical relative to hippocampal ripples (*N*=865). (**E**) Hippocampal relative to anterior thalamic ripples (*N*=81). Dashed error is 99% confidence interval of the null distribution. Data are from all channel pairs from all patients. Channel pairs with significant modulations only are depicted in Fig. 3. *P*-values computed using a Wilcoxon ranked-sum test to compare the modulation amplitude within -500 to 500 ms across bins and channel pairs for observed values versus null mean values. ns=non-significant, ****$p<0.0001$.

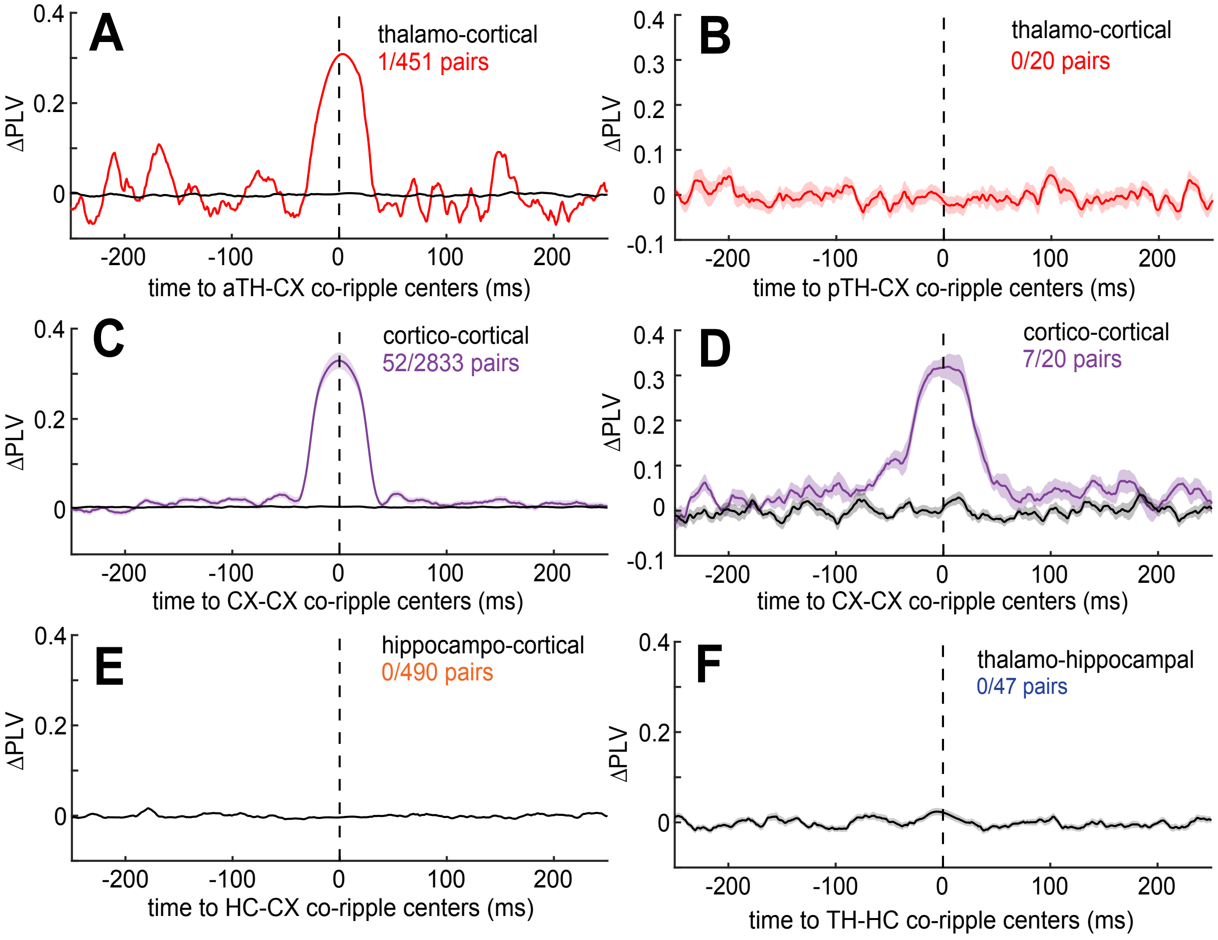


**Supplemental Figure 11. Thalamic ripples rarely phase-lock with cortical and never phase-lock with hippocampal ripples during NREM.** (**A-F**) Average and SEM ΔPLV time-courses across anterior thalamo-cortical (**A**, *N*=1/451 significant channel pairs, post-FDR $p<0.05$, randomization test), posterior thalamo-cortical (**B**, *N*=0/20), cortico-cortical (**C**, *N*=52/2833 from patients 1-10; and **D**, *N*=7/20 from patients 11-13), hippocampo-cortical (**E**, *N*=0/490), and thalamo-hippocampal (**F**, *N*=0/47). Time-courses in color show averages across significant and in black show averages across non-significant channel pairs. Channel pairs were only included if they had at least 40 co-occurring ripples with a minimum overlap of ≥25 ms. PLV, phase-locking value.

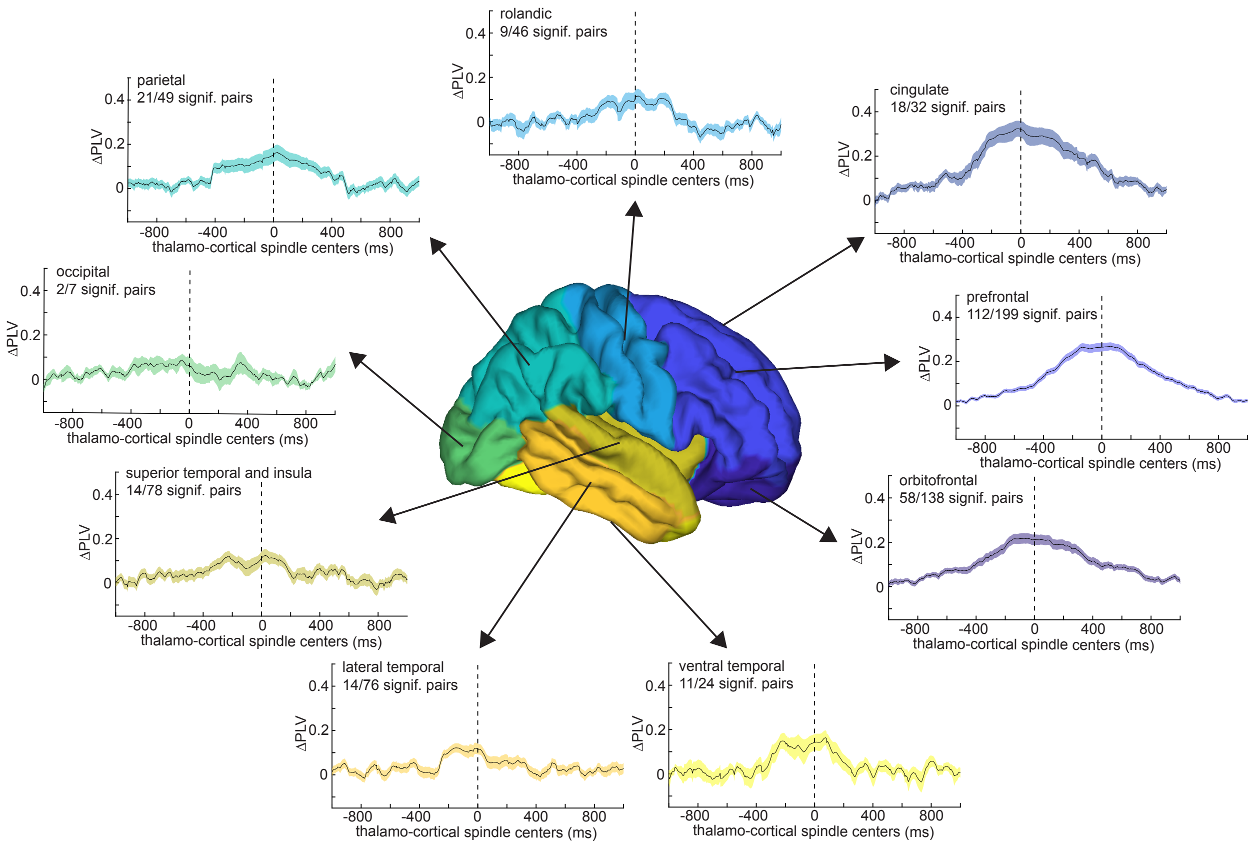


**Supplemental Figure 12. Thalamo-cortical spindle phase-locking results by cortical parcel.** Mean and standard error mean 10-16 Hz PLVs of anterior thalamo-cortical spindles across channel pairs. Proportions of significant thalamo-cortical channel pairs are indicated for each parcel. Note the greater proportion and magnitude of significantly phase-locked thalamo-cortical channels for anterior versus posterior cortical sites. Cortical parcels are amalgamations of the parcels in Desikan et al. (2006), as specified in the Methods.

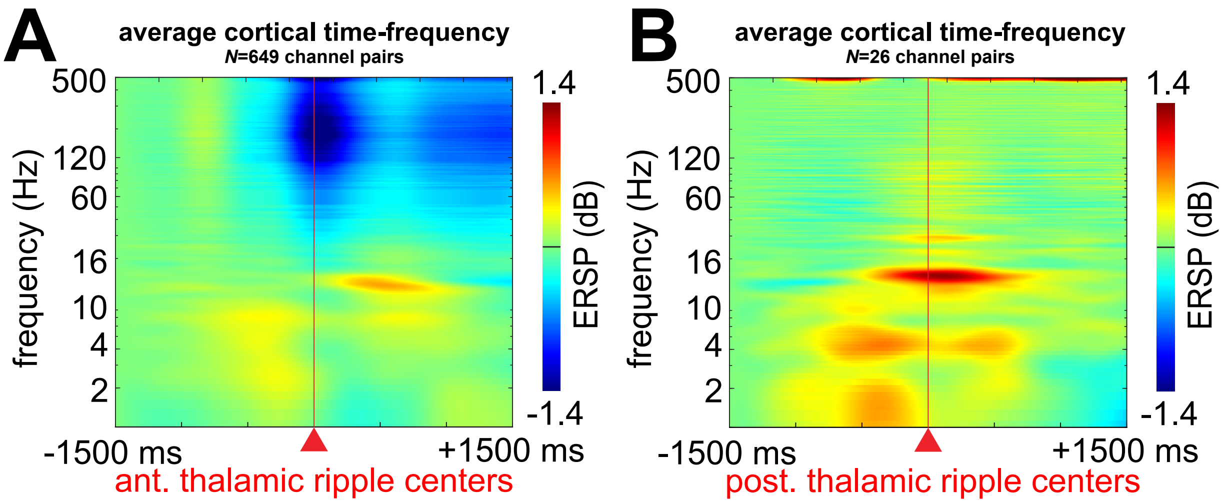


**Supplemental Figure 13. Cortical time-frequency locked to thalamic ripples.** (**A-B**) Average anterior (**A**) and posterior (**B**) thalamic-ripple locked time-frequency of cortical activity across thalamo-cortical channel pairs ($N_{aTH}$=649 and $N_{pTH}$=26).

| Pt. | Age | Sex | Hand. | Thalamic Ch. No. | Thalamic nuclei | Cortical Ch. No. | Hippocampal Ch. No. | Dur. (min) |
|---|---|---|---|---|---|---|---|---|
| 1 | 48 | F | L | 1R | Fa-Fa | 17R | 0 | 391.3 |
| 2 | 47 | F | R | 2R | AV-VA, VA-NRT | 28 (13L) | 10 (L: 3 pHC; R: 3 aHC, 4 pHC) | 128.9 |
| 3 | 43 | F | R | 1L | VLR-VLR | 22 (11L) | 2L pHC | 13.2 |
| 4 | 29 | M | R | 4L | AV-AV, AV-VA, VA-VLR, VLR-VLR | 28 (21L) | 3L (1 aHC, 2 pHC) | 282.5 |
| 5 | 33 | M | R | 2L | AV-VA, VA-VA | 47 (28L) | 2R (1 aHC, 1 pHC) | 92.9 |
| 6 | 61 | M | R | 3R | CM-VLR, VLR-VA, VA-WM | 19 (5L) | 2L pHC | 24.7 |
| 7 | 24 | M | R | 4R | AV-AV, AV-AV, AV-NRT, NRT-WM | 35 (14L) | 6 (R: 3 aHC, 2 pHC; L: 1 pHC) | 344.4 |
| 8 | 34 | M | R | 3R | VA-NRT, NRT-NRT, NRT-WM | 21R | 1R aHC | 77.7 |
| 9 | 52 | F | L | 2R | VA-VA, VA-NRT | 6R | 3R a HC | 562.2 |
| 10 | 38 | F | R | 2R | MDm-VLC, VLC-VLR | 38R | 2R pHC | 32.9 |
| 11 | 27 | F | R | 1R | PuM-PuM | 5R | 0 | 98.0 |
| 12 | 28 | F | R | 3R | PuM-PuM, PuM-PuL, PuL-PuL | 3R | 0 | 90.4 |
| 13 | 50 | F | R | 2R | PuM-PuL, PuL-NRT | 6R | 0 | 202.0 |

**Supplemental Table 1. Patient demographics and recording characteristics.** Channels are bipolar derivations included in the study. aHC, anterior hippocampus; AV, AnteroVentral; Ch., channel; CM, CentroMedian (of the ILTN- intralaminar nuclear complex); Dur., duration; Fa, Fasciculus (ILTN); Hand., handedness; No., number; MD, MedioDorsal; pHC, posterior hippocampus; Pt, patient; PuL, Lateral Pulvinar; PuM, Medial Pulvinar; NRT, Reticular nuc.; Pt., patient; VA, Ventral Anterior; VLR, Ventral Lateral, rostral division; VLC, Ventral Lateral, caudal division; WM, white matter. Maps of cortical channels are shown in Supplemental Fig. 2.

| Patient | Pre-implantation anti-epileptic drugs |
|---|---|
| 1 | Eslicarbazepine acetate, levetiracetam |
| 2 | Lamotrigine, zonisamide |
| 3 | Carbamazepine, Topiramate |
| 4 | Lacosamide, brivaracetam |
| 5 | Zonisamide, gabapentin, eslicarbazepine acetate |
| 6 | Carbamazepine, levetiracetam |
| 7 | Levetiracetam, lamotrigine |
| 8 | Lamotrigine, levetiracetam, clobazam |
| 9 | Brivaracetam, phenytoin |
| 10 | Lamotrigine, levetiracetam, gabapentin, carbamazepine |
| 11 | Unknown |
| 12 | Unknown |
| 13 | Unknown |

**Supplemental Table 2. Pre-implantation anti-epileptic drugs by patient.** Anti-epileptic drugs were tapered off or discontinued completely within 36-48 hours of admission to the epilepsy monitoring unit, where the recordings in this study were obtained. Long-acting medications, such as zonisamide, were held off prior to hospital admission preceding surgical implantation.

| Region | No. Ripples | Density ($min^{-1}$) | Amplitude (µV) | Frequency (Hz) | Duration (ms) |
|---|---|---|---|---|---|
| Anterior thalamus | 88320 | 19.8±4.6 | 1.79±0.91 | 92.4±0.6 | 62.6±5.9 |
| Posterior thalamus | 8996 | 11.3±2.2 | 1.95±0.34 | 90.2±0.4 | 93.3±23.0 |
| Cortex | 857420 | 21.9±2.5 | 4.17±1.69 | 90.1±0.5 | 65.3±3.6 |
| Hippocampus | 135646 | 21.3±4.7 | 11.39±7.23 | 88.5±1.3 | 74.8±7.9 |

**Supplemental Table 3. Ripple characteristics by region.** Values are counts or means and standard deviations across channels. See Fig. 1 for distributions and statistics.
